# Moss Transplants in the Tundra Reveal Host-Specific Microbiomes and Nitrogen Fixation Responses

**DOI:** 10.64898/2026.03.25.714306

**Authors:** Rebecca S. Key, Julia E. M. Stuart, Stuart F. McDaniel, Michael Hoffert, Emily Lockwood, Noah Fierer, Hannah Holland Moritz, Michelle C. Mack

## Abstract

In tundra ecosystems, moss-associated microbes are a major source of new nitrogen, yet the relative contributions of environment, host identity, and microbiome composition to variation in nitrogen-fixation rates are difficult to disentangle. To test how environmental change alters moss microbiomes and nitrogen-fixation rates, we used a one-year reciprocal transplant experiment between two Alaskan tundra sites that differ by 5°C in mean annual temperature. Intact moss cores containing one of three moss species, *Hylocomium splendens, Aulacomnium turgidum, and Pleurozium schreberi*, were transplanted between sites or returned to their home site. After one year, we quantified nitrogen-fixation rates using ^15^N incubation and characterized bacterial communities using 16S rRNA gene amplicon sequencing. *H. splendens* showed consistently low nitrogen-fixation rates with little transplant response, whereas *P. schreberi* and *A. turgidum* home and transplant tundra cores generally exhibited higher rates at the cooler, more northern site regardless of origin. In contrast, bacterial community structure changed little following transplantation, with composition driven primarily by moss species. Only in cyanobacteria and some heterotrophic bacterial lineages did we find subtle ASV-level changes. The absence of an association between microbial composition and nitrogen fixation, together with the heterogeneity among moss species, suggests that over short timescales, host physiology and microenvironment play a larger role in the variation of nitrogen-fixation rates than community turnover. The fact that short-term shifts in moss-associated nitrogen-fixation rate are driven primarily by host species identity, rather than microbiome restructuring, has important implications for near-term predictions of nitrogen inputs under Arctic climate change.

## Introduction

Mosses are key constituents of high latitude plant communities. In boreal and Arctic ecosystems, mosses account for 20-60% of net primary productivity on average (Chapin III *et al*. 1995, Bisbee *et al*. 2001, Turetsky *et al*. 2010). The presence of mosses affects carbon cycling both directly and indirectly through traits such as the production of recalcitrant litter and the insulation of soils (Cornelissen *et al*. 2007, Turetsky *et al*. 2012). Mosses also exert control over vascular plant growth through peat accumulation, soil temperature modulation, and allelopathy (Turetsky *et al*. 2012). Vascular plants in high latitude ecosystems are often nitrogen limited (Shaver and Jonasson 1999, LeBauer and Treseder 2008), with the largest source coming from nitrogen-fixing bacteria in association with moss hosts (Alexander and Schell 1973, DeLuca *et al*. 2002, Vile *et al*. 2014, Stuart *et al*. 2021). Mosses can readily access microbe-produced nitrogen while also limiting ecosystem nitrogen uptake in the soil via litter recalcitrance and high cation exchange capacity (Malmer *et al*. 2003, Cornelissen *et al*. 2007, Berg *et al*. 2013, Rousk *et al*. 2016), allowing them to compete effectively with vascular plants. Through these mechanisms, mosses have a disproportionate effect on carbon cycling dynamics in a region of the world that holds vast and vulnerable carbon stores (Lindo *et al*. 2013, Hugelius *et al*. 2014). Yet, the moss-associated nitrogen fixation rates vary tremendously at the micro-site level and across biomes. Identifying the drivers of this variation is critical to predict future nutrient fluxes in moss-dominated ecosystems as the climate warms.

Several lines of evidence show that both moss host identity and prevailing environmental conditions structure moss epihytic bacterial communities (Ininbergs *et al*. 2011, Bragina *et al*. 2012, Jean *et al*. 2020, Holland-Moritz *et al*. 2021). Mosses chemo-attract cyanobacteria (Bay *et al*. 2013) and exchange carbon and sulphur-containing compounds with their microbial associates in exchange for fixed nitrogen (Stuart *et al*. 2020). However, this interaction is not obligate as mosses transplanted from an area of low nitrogen deposition to an area of high nitrogen deposition experienced a rapid decline in cyanobacterial colonization (DeLuca *et al*. 2007). Transplant studies across other environmental gradients, such as nitrogen availability or plant community composition, have shown that microbial communities can respond quickly to changing conditions. For example, transplants of the mosses *Hylocomium splendens* and *Pleurozium schreberi* between birch and spruce stands resulted in slight microbiome shifts towards the new environment after transplantation (Jean *et al*. 2020). These findings suggest that while mosses recruit a distinct, species-specific bacterial consortia, overall community composition is dynamically regulated by moss or microbes in response to environmental conditions.

Rates of moss-associated microbial nitrogen fixation depend on the presence and activity of diazotrophic bacteria, which in turn may be influenced by the moss host species (Stuart *et al*. 2021) and environmental conditions like temperature, moisture, and light availability (Gundale *et al*. 2012a; Rousk *et al*. 2017; Rousk & Michelsen, 2017). Mosses differ in shade tolerance, cold adaptations, microhabitat preference, and water retention (Mills and Macdonald 2004, Elumeeva *et al*. 2011, Jonsson *et al*. 2015), which can collectively result in variable responses of nitrogen-fixation rates to changes in conditions across moss species (Leppänen *et al*. 2015, Stuart *et al*. 2021). Short-term increases in temperature can increase nitrogen-fixation rates, but long-term *in situ* warming experiments found either no effect or a decrease in moss-microbe nitrogen fixation (Gundale *et al*. 2012a; Sorensen *et al*. 2012; Permin *et al*. 2022). Bacteria with alternative forms of nitrogenase using vanadium or iron are likely less sensitive to low temperatures than molybdenum nitrogenases, and moss-associated N₂ fixation has been observed year-round, even showing peaking activity outside of summer months (Darnajoux *et al*. 2017, Bellenger *et al*. 2020, Andersen *et al*. 2025). Evaluating the interactions between environmental conditions and host moss identity is crucial, as both can independently or collectively influence microbial nitrogen inputs in tundra ecosystems, especially under changing climatic conditions.

To test the influence of environmental factors in combination with host species identify on moss-associated microbial communities and nitrogen-fixation rates, we conducted a reciprocal transplant experiment between tundra in Healy, Alaska, and Toolik Lake Field Station. We surveyed three common tundra moss species (*Aulacomnium turgidum, H. splendens, and P. schreberi*) using 16S rRNA gene sequencing and nitrogen-fixation measurements, and recorded the neighboring plant community. We hypothesized that if nitrogen-fixation rates changed in transplants without concurrent shifts in microbial community composition, this would suggest that variation in nitrogen-fixation changes was driven by moss host. In contrast, if changes in nitrogen fixation coincide with shifts in particular microbes, this would suggest an environmentally mediated microbial contribution to nitrogen fixation. Given the strong influence of host identity on moss-associated microbiomes, we further expected host-specific differences in microbial composition and nitrogen-fixation responses, as well as spatial differentiation in microbial communities between the southern and northern site. Alternatively, if neither nitrogen-fixation rates nor microbial community composition differ between mosses remaining in their home environments and those transplanted to new sites, this would suggest that moss–microbe associations are relatively robust to the environmental variation examined in this study.

## Materials and Methods

### Study Sites and Experimental Design

In July 2018, we selected two moist acidic tundra sites in Healy near Eight Mile Lake (65°52′51″ N, 149°14′12″ W) and Toolik Lake Field Station (68°37′27″ N, 149°36′18″ W), Alaska, USA (Figure 1). Sites were similar in elevation and mean annual precipitation (MAP), and not similar in flora and mean annual temperatures (MAT). The MAT at Toolik is -6.4 °C and -1.0 °C at Healy. Both sites were near, but not within, long-term experimental plots.

**Figure 1.**
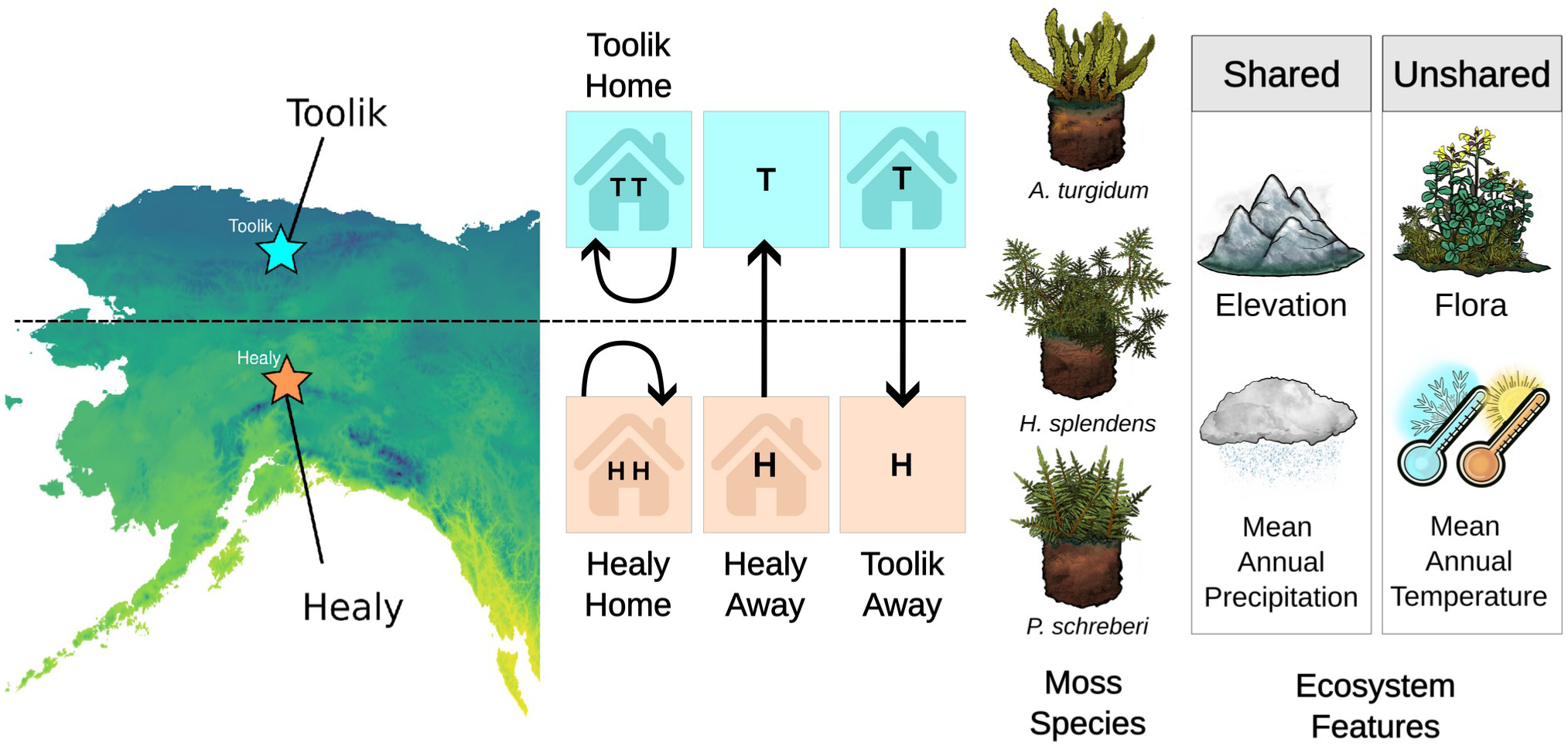
Experimental Design and Study Sites. (**Left**) A temperature gradient map of Alaska showing two study sites: the northern field station located in Toolik (68°37’27”N, 149°36’18”W) and the southern field station located in Healy near Eight Mile Lake (65°52’51”N, 149°14’12”W). Darker colors indicate lower mean annual temperatures (MAT) while lighter colors indicate higher MAT. (**Center**) Schematic of the reciprocal transplant design. At each site, moss cores (30 cm diameter, ∼15 cm depth) were excavated and randomly assigned to two treatment groups: “Home” (Toolik Home, TT; Healy Home, HH) or “Away” (Toolik Away, TH; Healy Away, HT), with Away cores transplanted between sites within 48 hours. The three target moss species include *Hylocomium splendens, Pleurozium schreberi,* and *Aulacomnium turgidum*. Plant communities also present in each core were surveyed at the time of transplantation. After one year, cores were revisited to re-survey plant communities and collect moss tissue samples for 16S sequencing and nitrogen fixation measurements. (**Right**) Both sites share elevation and mean annual precipitation (MAP), but differed in surrounding plant communities and MAT (Toolik: −6.4°C, Healy: −1.0°C).

At each site, we excavated twelve circular cores randomly from areas where three moss species were observed: *Hylocomium splendens (Hedw.) Brid., Pleurozium schreberi (Brid.) Mitt., and Aulacomnium turgidum (Wahlenb.) Schwgr.* Each core was 30 cm in diameter and approximately 15 cm in depth. We selected patches from inter-tussock spaces, with at least one meter in between core edges. Six of the twelve cores were designated as the home (*i.e.*, untransplanted) treatment, the other as the away (*i.e.*, transplanted) treatment. We immediately replaced the away cores with the home cores, transferred the away cores in buckets containing multiple cold packs, stored these buckets in refrigerated rooms overnight, and transported them to their new site. While transporting the away cores, efforts were made to keep cores intact and cool. All transplantations were completed within 48 h.

Following transplantation into either home or away environments, other plants which were present on the core were surveyed by counting the number of vascular species (*e.g.,* angiosperms) and taking visual estimates of non-vascular (*e.g.,* mosses, liverworts, hornworts) and lichen species to obtain a percent cover. Mosses were identified by genus- or species-level when possible, while liverworts and lichens were identified to genus if cover exceeded 5% in a core. We did not measure *Sphagnum* spp., other acrocarpous species, or feather mosses. We re-assessed each core again at the end of experimentation in August 2019, before taking nitrogen-fixation measurements.

### Weather Data Collection

MAT and PAR data for the Toolik site were provided by the Toolik Lake Field Station Environmental Data Center (Environmental Data Center Team, 2021), while data for Healy were provided by the Schuur laboratory through the Bonanza Creek LTER (Schaedel *et al*. 2021). MAT and PAR datasets were checked for anomalies in the time leading up to nitrogen fixation measurements. Spatial temperature data for the state of Alaska was sourced from the ClimateNA database (Wang *et al*. 2016). The historical normal annual dataset for the period 1991-2020 was used to generate a temperature gradient map, spanning longitudes -170° to -130° W and latitudes 51° to 71° N.

### Nitrogen Fixation Measurements

One year after transplantation, we measured nitrogen-fixation rates for each core by first removing 10 moss ramets of a single species and placing them in an airtight 60 mL polypropylene syringe. Each moss ramet was approximately 5 cm of length, cleaned of debris and other plant material, and contained both green and senescent moss tissue. Ramets were lightly sprayed with distilled water to avoid desiccation during incubation. Within hours of initial collection, we plunged 10 mL of ambient air into each syringe before adding 10ml of 98 at% enriched ^15^N_2_ gas, making the final volume 20 mL (Sigma-Aldrich Inc., Lot MBBB9003V). We then incubated for 24 h under ambient light conditions. Afterwards, moss samples were dried at 60°C for 48 h and shipped to Northern Arizona University. Samples were homogenized by finely chopping with scissors and rolling 6 mg of tissue into a tin capsule. These were then run on a Costech ECS4010 elemental analyzer coupled to a Thermo Scientific Delta V Advantage isotope ratio mass spectrometer to obtain percent nitrogen, percent carbon, δ^15^N, and δ^13^C measurements.

We calculated atom percent enrichment (APE) for each sample by subtracting the natural abundance atom percent from the incubated sample. Natural abundance was estimated using species-specific mean δ¹⁵N values (*A. turgidum* = −3.393‰, *P. schreberi* = −3.598‰, *H. splendens* = −3.343‰) from Stuart *et al*. (2020). We then scaled isotopic uptake by the sample weight and the air:tracer ratio to calculate total (^15^N+^14^N) nitrogen fixation (Jean *et al*. 2018). Rates are expressed as µg N g⁻¹ moss day⁻¹. The average enriched δ¹⁵N value averaged across all samples was 2573.3 ± 148.4. For additional details on nitrogen fixation methodology, see Stuart *et al*. 2020.

### Amplicon Collection, Sequencing, and Processing

Dried moss samples were shipped to the University of Florida. Genomic DNA was extracted using the MoBio Power Soil DNA extraction kit (MoBio Laboratories, Carlsbad, CA) on 0.25 g of homogenized frozen tissue. DNA was PCR amplified in triplicate using the 515F/806R primer pair spanning the V4 16S rRNA gene. Amplified products were bar coded, normalized to equimolar concentrations using ThermoFisher Scientific SequalPrep Normalization plates (Thermo Fisher Scientific Inc. USA), pooled, and sequenced using 150-bp paired-end reads on the Illumina Miseq platform at the Center for Microbial Exploration at the University of Colorado, Boulder.

Amplicon sequence variants (ASVs) were resolved using DADA2 denoising (Callahan *et al*. 2016) implemented in Bioconductor v3.10. MiSeq FASTQ files were demultiplexed using Idemp. Reads containing uncalled bases and shorter than 50 bp were removed. Adapter sequencing were removed with Cutadapt v4.1 (Martin, 2011). DADA2 filtering parameters were truncLen = c(150,150), maxEE = c(2,2), truncQ = 2, maxN = 0, rm.phix = TRUE, retaining only 150 bp paired-end reads with ≤ 2 expected errors. These error rates were learned from truncated reads using 10 million bases in randomized mode. ASVs were inferred in pooled mode, merged across forward and reverse reads, and chimeras were removed using the consensus method. ASV taxonomy was assigned using the SILVA 16S reference database v132 (Quast *et al*. 2013). Both mitochondria and chloroplast ASVs (2.6% and 6.39% of all ASVs, respectively) were discarded before subsequent analysis. Negative control (blank) samples showed minimal contamination, with ≤50 reads per sample on average and no shared ASVs.

### Linear Mixed-Effects Modeling

Statistical analyses were performed in RStudio v1.2.1335 using R version v3.6.1 (R Core Team, 2019). Data organization and visualization were performed using tidyr, dplyr, and ggplot2 (Wickham *et al*. 2019). To test the hypotheses of this study, we used linear models and linear mixed-effects models in lme v4 1.1-21 (Bates *et al*. 2015) paired with Satterthwaite’s degrees of freedom method in lmerTest (Kuznetsova *et al*. 2017). We inspected all model residuals visually using the check_model() function in the performance package and displayed model results through the ggeffects package (Lüdecke 2018, Lüdecke et al. 2021). Additional analyses used the vegan v2.5-7 package for nonmetric multidimensional scaling, running permutational multivariate analysis of variance (PERMANOVA) tests on community composition data, and creating treatment group ellipses (Oksanen *et al*. 2025).

Linear mixed-effects modeling was performed to test the effect of transplantation on moss-associated nitrogen-fixation rates. The rate of nitrogen fixation was treated as a response variable, while the interaction between home site (Toolik or Healy) and transplant status (home or away) was treated as fixed effects. Moss species were treated as a random effect due to the assumption that moss species will affect the results but could not be directly tested in this model due to unequal distribution between treatment groups. Sampling core was also treated as a random effect to account for autocorrelation within cores but removed it from the final model as the between-group variability was not sufficient to warrant its incorporation.

Given the constraints of sample size, species-specific responses to transplantation were evaluated using separate linear models. For these models, core was not included as a random effect as only one sample of each species was measured in each core. Since *A. turgidum* was well represented in all treatments, nitrogen-fixation rates associated with those samples were regressed against an interaction between home site and transplant status. For *H. splendens* and *P. schreberi*, only samples originating in Toolik or Healy, respectively, were included in the models. For each species model, nitrogen-fixation rates were regressed with transplant status to compare mosses that remained in their home environment to ones that were transported to a novel environment.

### Microbe Community Analysis

Microbial community analyses used vegan v.2.6-4 (Oksanen *et al*. 2025), phyloseq v1.44.0 (McMurdie and Holmes 2013, 2014), tidyverse v2.0.0 (Wickham *et al*. 2019), and ANCOM-BC v.2.6.0 (Lin and Peddada 2020) packages. ASVs were first rarefied at a threshold of 5,400 reads using rarefy_even_depth(rngseed = 1, replace = FALSE). Second, relative percent abundances was calculated by dividing ASV count observations by the chosen rarefy threshold (*i.e.*, the number of observations per sample). Observed, Shannon, and Simpson diversity metrics were calculated using estimate_richness(). These values were compared to non-rarefied diversity values before using the rarefied diversity values to compare home and away treatment groups. Statistical significance and effect sizes were calculated using ANOVA followed by Cohen’s d and Tukey’s HSD tests. Using sample ID as rows and ASVs as columns, both Principal Coordinates Analysis (PCoA) using Bray-Curtis dissimilarity and hierarchial clustering was performed using phyloseq. Post-hoc fitting of categorical variables (*e.g.*, moss species, treatment group) and continuous variables (*e.g.,* nitrogen fixation measurements) was performed using the ‘envfit’ function. Differential abundance analysis was conducted using ANCOM-BC package. Samples were stratified into three contrast groups: (1) Healy home vs. Healy away, (2) Toolik home vs. Toolik away, and (3) Healy home vs. Toolik home, across each moss species. Only ASVs present in more than 10% of samples (prv_cut = 0.10) were used. P-values were adjusted using Benjamini-Hochberg correction (p_adj_method = “BH”) to account for multiple group comparisons. To avoid uneven sample distributions, only comparisons with at least three replicates per moss species were evaluated. Significant ASVs with adjusted P < 0.05 were considered significant and classified as either enriched (positive fold change) or reduced (negative fold change). Phylum-level composition was summarized by tallying the number of significant ASVs.

### Plant Community Analysis

To assess changes in the plant community composition, we generated non-metric multidimensional scaling (NMDS) ordinations using the Bray-Curtis dissimilarity index separately for the vascular and non-vascular community composition of each core in 2019. Each ordination used two dimensions after confirming a reduction in stress (< 0.05) from adding additional axes. After checking for homogeneity of group variances, we ran a PERMANOVA for each community to test for the effect of the interaction between home site and transplant status using the ‘adonis2’ function with the Bray-Curtis dissimilarity index and 999 permutations. Depending on PERMANOVA results, 95% confidence interval ellipses were constructed in ordination space for each group and then extracted for plotting. Using the function ‘envfit’ we also fit species vectors within each community.

## Results

### Nitrogen Fixation

Nitrogen-fixation rates within the transplant experiment were affected by the interaction between home site and transplant status when host species was treated as a random effect (Figure 2, F = 13.18, P < 0.001). Mosses from the Healy Home group (Healy to Healy) fixed nitrogen at lower rates compared to the three other groups, while mosses from the Healy Away group (Healy to Toolik) fixed as much nitrogen as mosses from the Toolik Home group (Toolik to Toolik) (Figure 2). The random effect of moss species accounted for 64% of variance explained within the model. The marginal and conditional R^2^ of model was 0.11 and 0.68, respectively.

**Figure 2.**
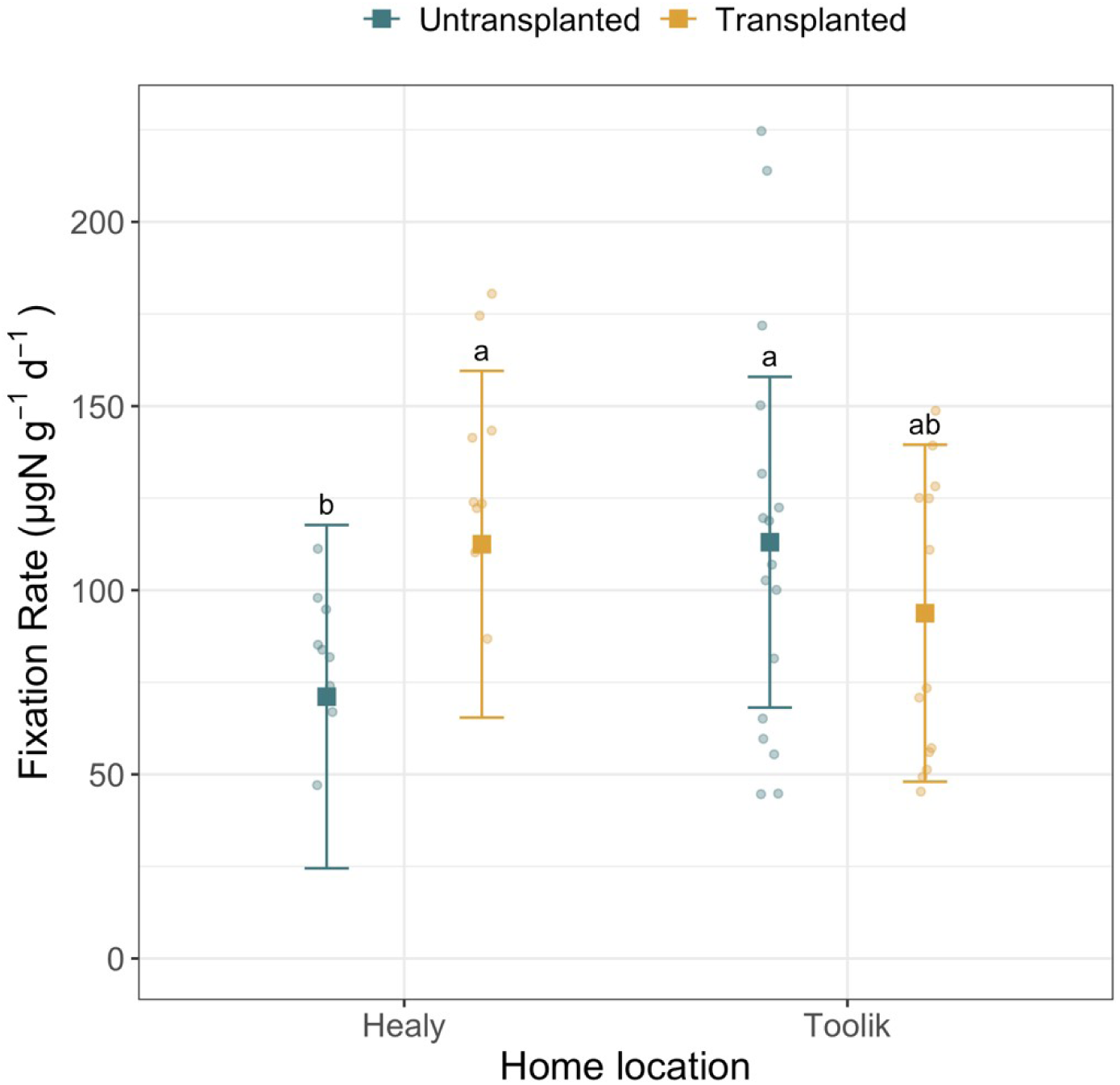
Nitrogen Fixation Rates Across Moss Transplant Groups. Model-predicted means (squares) and standard deviations (error bars) as well as raw data (round points) for nitrogen fixation rates, controlling for moss species (random effect). Home sites are on the x-axis, and colors represent whether a sample was transplanted to the alternate site (yellow) or re-transplanted back to the home site (blue). The two outside groups, far left and far right, were measured in Healy, while the inner two inner groups were measured in Toolik as the final site.

Nitrogen-fixation rates from each moss species showed distinct responses to transplantation. *H. splendens* did not respond to transplantation, showing no change between Toolik Home and Toolik Away groups, and it showed the lowest fixation rates compared to the other two moss species (Figure 3, F = 0.28, P = 0.611). *A .turgidum* was present in almost all cores, and its associated nitrogen-fixation rates were modeled as an interaction between home site and transplant status. For *A. turgidum,* both home groups had the strongest impact on nitrogen-fixation rates, with Toolik Home having higher rates than Healy Home (F = 4.79, P = 0.04). However, the interaction between home sites and transplant status appeared to have a moderate effect compared to transplant status by itself (F = 3.71, P = 0.072 for interaction term, F = 0.84, P = 0.374 for transplant status). Lower nitrogen fixation rates were observed for both Healy and Toolik Away groups compared to the Toolik Home group, although both away groups fixed more nitrogen than the Healy Home group. *P. schreberi* responded strongly to transplantation, with the Healy Home group showing lower nitrogen fixation rates than the Healy Away group (Figure 3, F = 42.52, P < 0.001).

**Figure 3.**
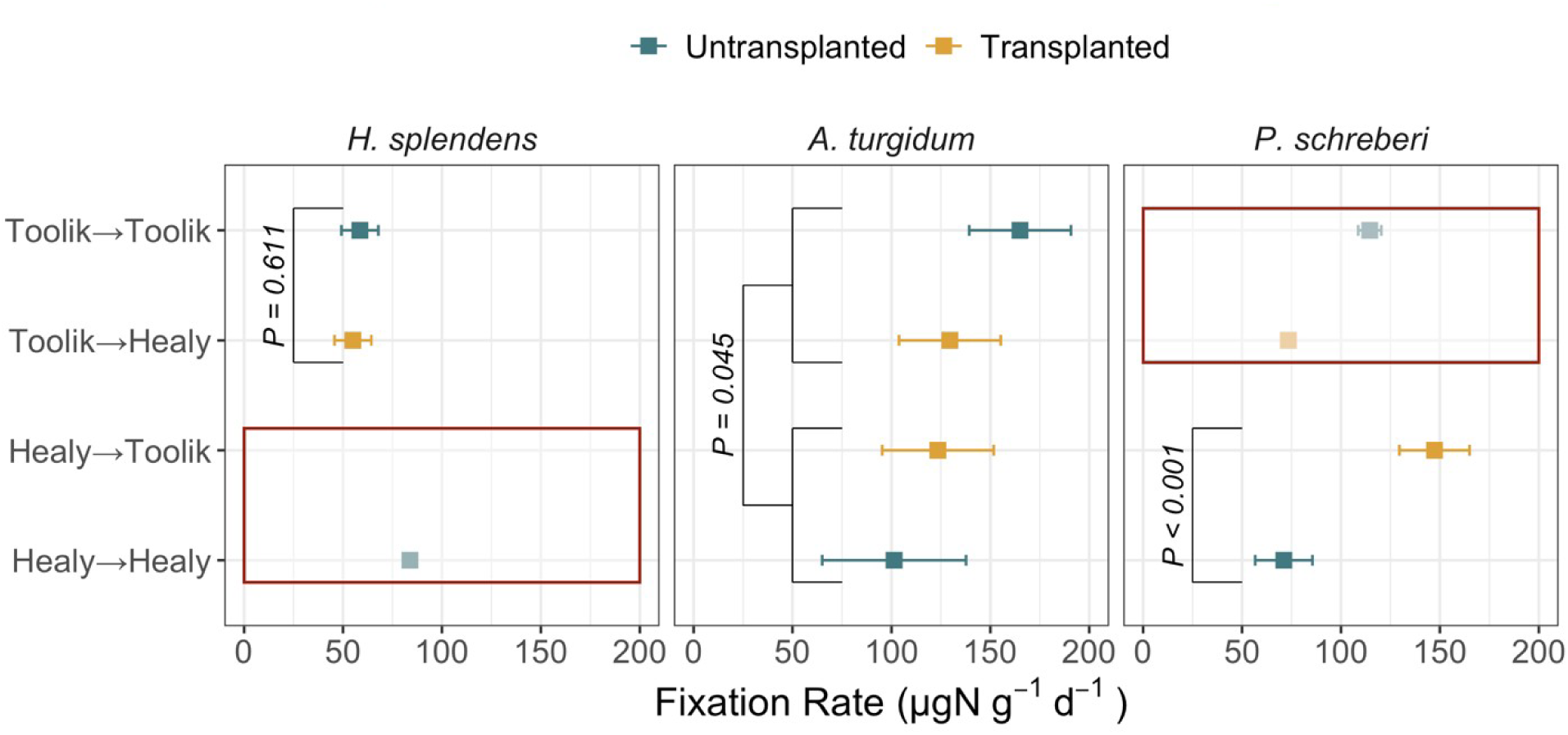
Distinct Responses of Moss-Associated Nitrogen Fixation to Transplantation. Individual linear models were run for each tested moss species, and each model here is represented in one box. The *H. splendens* model included only samples from Toolik, and the *P. schreberi* model contained only samples from Healy due to incomplete replication within treatment groups. Red boxes were placed over data excluded from the models and within those areas squares represent group mean and bars are standard error. Otherwise, points represent estimated marginal means and bars are 95% confidence intervals from models. *H. splendens* did not respond to transplantation, while *P. schreberi* responded strongly. *A. turgidum* nitrogen fixation rates were most affected by home site but there was a moderate interaction between home site and transplant status (P=0.07).

### Moss Microbe Communities

We examined composition differences of 16S ASVs across treatment groups (Figure 4). The 16S dataset consisted of 36 samples: 15 derived from *A. turgidum*, 12 from *P. schreberi*, and 9 from *H. splendens* (Figure S1). A total of 5,905 ASVs were captured across samples after rarefying: 2,201 ASVs from *A. turgidum*, 1,745 from *P. schreberi,* and 1,958 from *H. splendens* (Figure S2). 908 ASVs were shared across the three species. PCoA ordination revealed separation of samples primarily driven by moss species (F = 2.9051, P = 0.001, R² = 0.1416), followed by home site (F = 2.9650, P = 0.001, R² = 0.0723) (Figure 4A; Figure S3). Transplant condition and treatment group accounted for 2.5% (F = 1.0326, P = 0.382) and 5.4% (F = 2.2275, P = 0.001) of the variance, respectively, though the former was not statistically significant (Figure S3).

**Figure 4.**
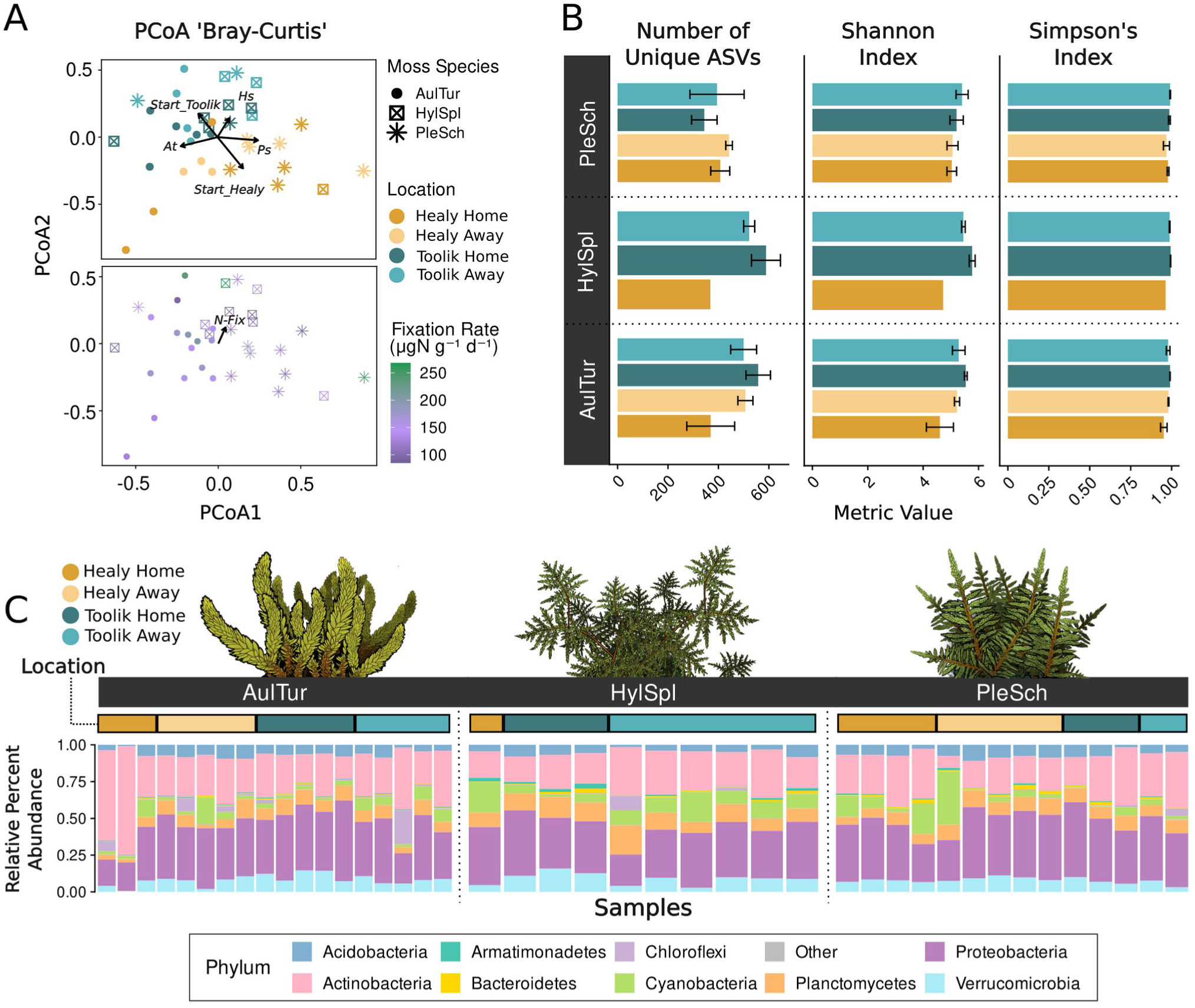
Moss Microbial Diversity and Composition **(A)** Non-metric Multidimensional Scaling (NMDS) with Bray-Curtis dissimilarity. The upper NMDS plot shows the sample ordination, with points colored by treatment group and shaped by moss species. Arrows represent categorical vectors, showing the influence of moss species and home site on 16S community composition. Moss species include: *Aulacomnium turgidum* (AulTur; At), *Hylocomium splendens* (HylSpl; Hs), and *Pleurozium schreberi* (PleSch; Ps). ‘Start_H’ means all samples originated from Healy, while ‘Start_T’ means those originating from Toolik. The lower NMDS plot displays the same ordination but with points colored by nitrogen fixation rate (N-Fix). The arrow represents the continuous vector and its influence on the ordination space. **(B)** Bar charts show the average values of three diversity metrics: Observed number of ASVs (richness), Shannon Index, and Simpson Index for each treatment group. Moss names and treatment group colors are the same as Panel A. The x-axis shows diversity values for each metric, while the y-axis shows each group category. **(C)** Bar plots display relative percent abundance at the phylum-level across samples. Each bar is an individual sample, colored by Phylum. The feature bar above each plot is color-coded by treatment group. Samples are grouped by moss species and site.

For Observed (*i.e.,* the number of unique ASVs), Shannon-Weaver, and Simpson diversity metrics, minimal and non-significant changes were observed (Figure 4B; Figure S4), with the exception of Toolik Home and Healy Home groups for *A. turgidum* (P = 0.042; Cohen’s d = 1.91). Notably, diversity metrics also remained unchanged before and after rarefying samples (Figure S5). For each moss host, the majority of ASV relative percent abundance belonged to four dominant phyla: Proteobacteria (30-35%), Actinobacteria (24-31%), Planctomycetes (7-11%), and Verrucomicrobia (7-9%) (Figure S6). This dominance profile remained consistent across moss species, with exception of Cyanobacteria communities in *H. splendens* which contained higher relative percent abundance than Verrucomicrobia. Hierarchical clustering at phylum-level revealed that sample dissimilarity was driven by home site, with Toolik samples appearing to cluster for each moss species (Figure S7). Examining phylum-level percent abundance changes across samples, there was no observable differences across moss species or treatment groups (Figure 4C).

To identify ASV-level changes in response to transplantation, we performed an ANCOM-BC differential abundance analysis (Figure 5). Due to insufficient replication in *H. splendens*, we applied tests only to *A. turgidum* and *P. schreberi* (Figure 5A). Three contrasts were tested: a Healy Transplant contrast (Healy Home vs Healy Away), a Toolik Transplant contrast (Toolik Home vs Toolik Away), and a Home Control contrast (Healy Home vs Toolik Home). For each moss species, the Home Control contrast identified ASVs that differed between the two home sites, reflecting taxa structured by spatial site differences rather than by transplantation, whereas both transplant contrasts identified ASVs whose abundances shifted under non-native site conditions. Categorizing ASVs based on overlap among these three contrasts yield three distinct scenarios (Figure 5B). The first scenario included site-specific ASVs that differed between home sites. The second scenario included ASVs overlapping between the Home Control and Transplant contrasts, representing taxa that differ between sites and change in relative percent abundance following transplantation. The third scenario included ASVs present in the Transplant contrasts but absent from the Home Control, representing taxa that do not differ under home conditions yet change in relative percent abundance following transplanted.

**Figure 5.**
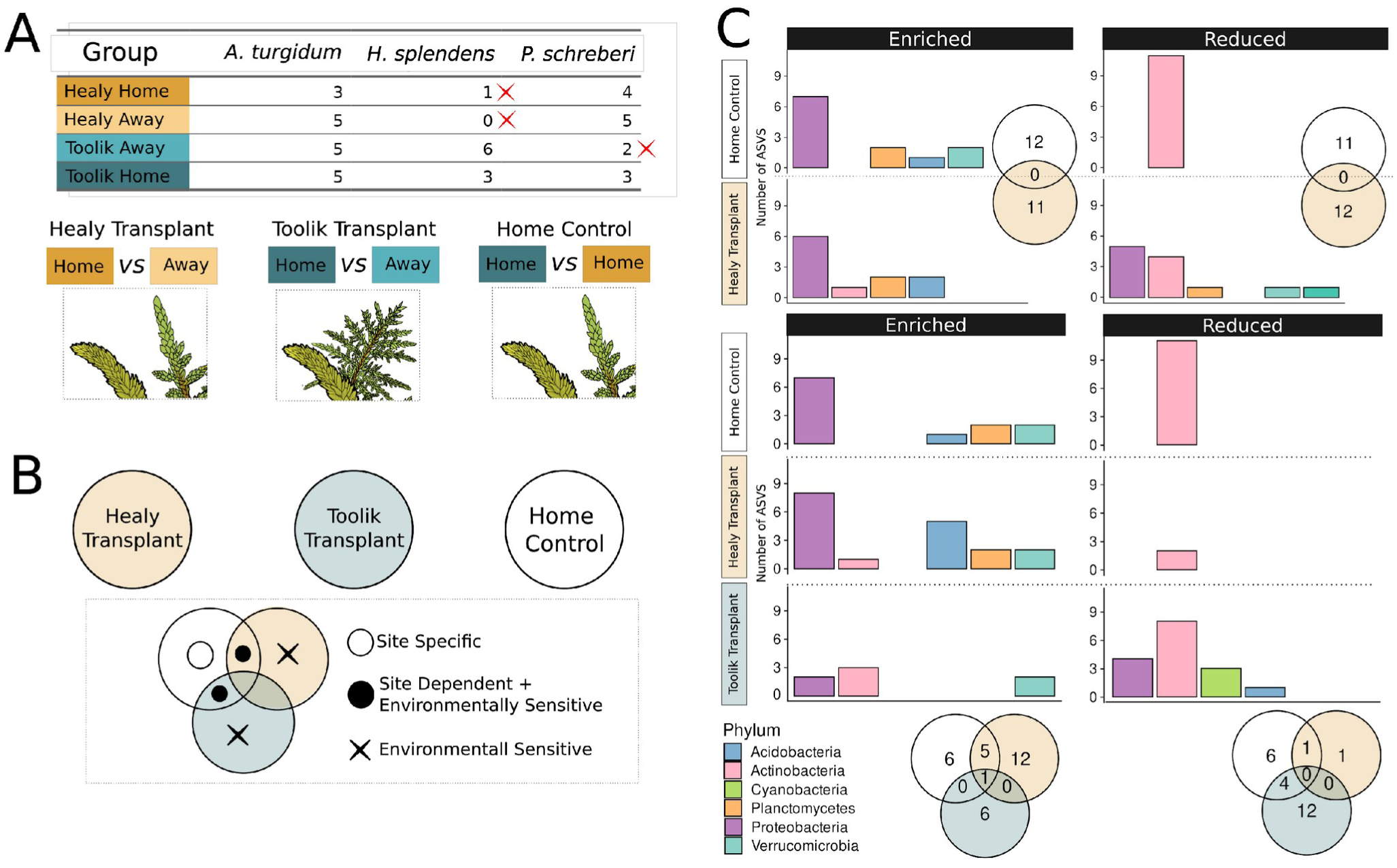
ASV-Level Shifts in Moss Microbial Communities **(A)** ANCOM-BC Test Design and Sample Distribution. The table shows the number of samples per moss species across four treatment groups: Healy Home, Healy Away, Toolik Home, and Toolik Away. Red Xs denote groups with fewer than three replicates. These treatment were combined into three contrasts: Healy Transplant, Toolik Transplant, and a Home Control. Only moss species with sufficient replication in the Home Control and at least one Transplant contrast were included in ANCOM-BC. **(B)** Venn diagram illustrating how taxa were categorized from ANCOM-BC results. White circles (ASVs changing between Home sites only) represent site-specific taxa not affected by transplantation. Black circles (ASVs overlapping Transplant and Home Control comparisons) indicate site-dependent, environmentally responsive taxa. X symbols (ASVs present in both Transplant contrasts but absent in the Home Control) mark taxa that responded to a transplant-associated factor independent of site identity. **(C)** Bar plots showing the number and phylum-level identity of enriched and reduced ASVs across Home Control, Healy Transplant, and Toolik Transplant comparisons for each moss species. Venn diagrams summarize the overlap of enriched and reduced ASVs across the two transplant sites and home conditions, indicating whether responses are shared or site-specific. Colors correspond to major bacterial phyla contributing to each response category.

For *A. turgidum*, we found 30 significantly enriched and 24 significantly reduced ASVs across the three contrast groups (Figure 5C; bottom Venn diagrams). Focusing on third scenario ASVs in the Toolik Transplant contrast with relative percent abundance shifts exceeding 1 log_2_ fold change, we observed Cyanobacteria, Proteobacteria, and Acidobacteria ASVs that were stable between home conditions yet responded to transplantation. Notably, cyanobacterial ASVs such as *Nostoc* PCC-73102 (P = 0.048), *Stigonema* SAG 48.90 (P = 0.04), and an unclassified Nostocaceae member (P = 0.02) were detected, along with other significant ASVs from Solirubrobacteraceae, Frankiaceae, Burkholderiaceae, and Acetobacteraceae (Figure 5C; Table S1). For *P. schreberi*, 23 ASVs were enriched and 23 were reduced (Figure 1C; upper Venn diagrams). Under the third scenario and log_2_ fold change condition, notable ASVs included two Planctomycetes ASVs belonging to *Singulisphaera (*P = 0.009) and the WD2101 soil group (P = 0.01), one Verrucomicrobia ASV assigned to Chthoniobacteraceae (P = 0.01), and one unclassified ASV from Armatimonadales (P = 0.04) (Figure 5C; Table S2).

### Plant Communities and Weather Trends

Neither the vascular nor the non-vascular plant community responded to transplantation or the interaction between home site and transplant status (Vascular community: F = 1.02, P = 0.412 for transplant status; F = 1.37, P = 0.240 for interaction term; Non-vascular community: F = 0.46, P = 0.680 for transplant status; F = 0.66, P = 0.482 for interaction term). Conversely, for both plant community types, home site was significant (Figure 6; Vascular community: F = 6.29, P < 0.001; Non-vascular community: F = 22.95, P < 0.001). Within the non-vascular community species vectors, the three target moss species had a P value smaller than 0.01 (Figure S9). Several vascular species, including *Rubus chamaemorus, Rhododendron tomentosum, and Cassiope tetragona,* had low P values and appeared to be important in structuring differences between communities (Figure S10).

**Figure 6.**
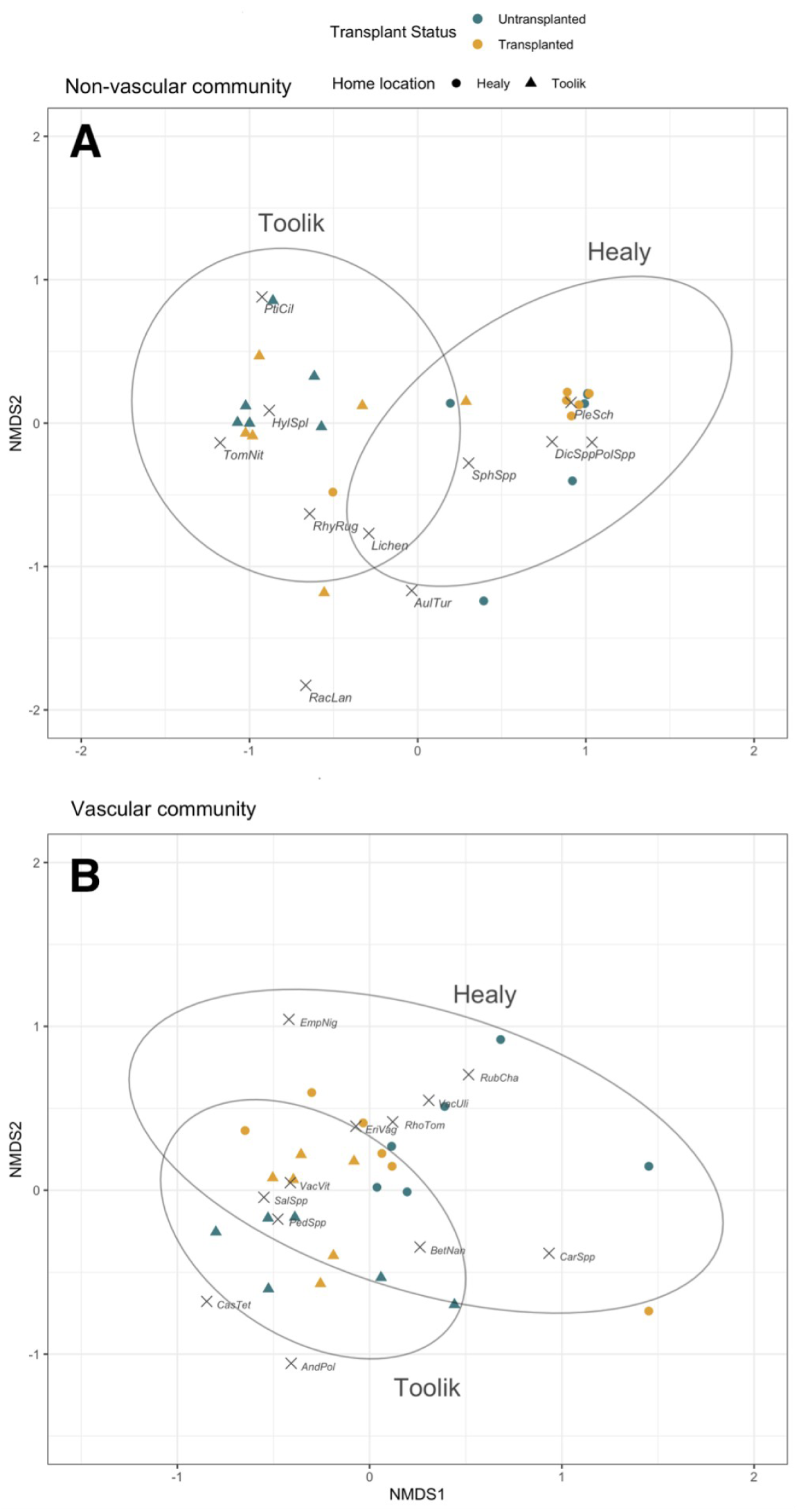
NMDS Ordination of Observed Non-Vascular and Vascular Plant Communities NMDS ordination plots of plant community structure. Ellipses representing the 95% group confidence interval using the standard deviation of points for home location in the non-vascular (A; upper) and vascular (B; lower) plant communities. perMANOVA results showed no effect of transplant status. Species ordination scores are represented by X’s. Species codes are the first three letters of the genus followed by the first three letters of the species. If species were not differentiated within a genus, *Spp* appears in place of the species code.

In the days leading up to the nitrogen incubations, average daily PAR at Toolik was higher compared to Healy (Figure 7). Healy had more precipitation in the summer of 2019, with 297 mm total falling during the two-month period prior to incubation. Out of those 60 days, 34 included some precipitation falling. In Toolik, the same two-month period before incubation included 208 mm of rain and 33 rainy days. In the fortnight preceding nitrogen fixation measurements, Toolik received 73 mm of precipitation to Healy’s 158 mm, with 10 and 11 days of rain events, respectively. As expected, daily mean and maximum temperatures were higher in Healy compared to Toolik. Relative humidity was higher in Healy ( >75% in all days leading up to incubation) than at Toolik (50-80%, Figure S11).

**Figure 7.**
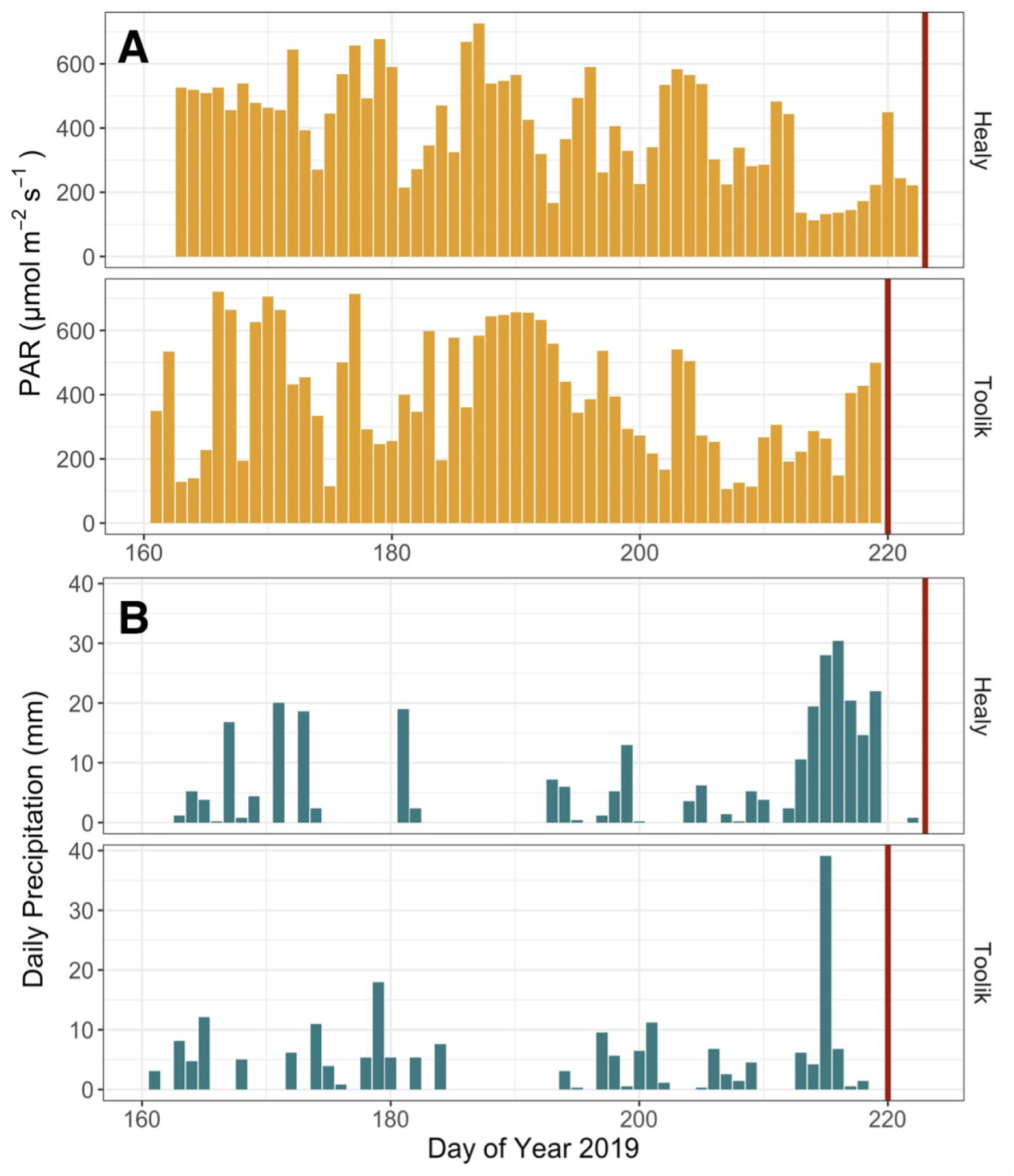
Environmental Patterns Leading Up to Nitrogen Fixation Measurements Bar charts showing average daily photosynthetically active radiation (PAR) (A) and precipitation (B) in the growing season leading up to nitrogen fixation measurements. Dates of incubation are indicated by a red bar on each graph. The *x*-axis is the Julian day of year.

## Discussion

It is widely known that moss-associated microbes are a major source of new nitrogen in tundra ecosystems, yet the relative contributions of host identity, microbiome composition, and environment to variation in nitrogen-fixation rates are difficult to disentangle. Using a reciprocal transplant between tundra at Toolik Lake Field Station and Healy, Alaska, we found species-specific functional responses to transplantation in moss-associated nitrogen fixation that were independent of microbial and neighboring plant community composition. Nitrogen-fixation rates were higher for mosses located at Toolik, regardless of whether they were native or transplanted from Healy (Figure 2). However, the magnitude and direction of this response varied among moss species (Figure 3). In contrast, the microbial composition at the taxonomic resolution of 16S amplicons remained stable following transplantation (Figure 4B and 4C), with only subtle ASV-level changes (Figure 5C). Below we examine the factors that may contribute to the host species-specific response in nitrogen fixation, and the potential consequences of this variation.

### Species-Specific Responses for Nitrogen Fixation

Overall, the nitrogen-fixation rates we observed fall within the range previously reported for moss-associated fixation in Alaska and Northern Europe (Bay *et al*. 2013, Vile *et al*. 2014, Jean *et al*. 2018). For *P. schreberi*, transplantation to the Toolik site resulted in nearly twofold higher nitrogen-fixation rates compared to the untransplanted Healy control (Figure 3). By comparison, *H. splendens*, a closely related feathermoss, displayed lower rates with little evidence of phenotypic plasticity. *A. turgidum* showed a site-specific profile, with higher rates at the northern site in untransplanted samples, suggesting potential genetic differences between southern and northern populations. Together, these patterns indicate heterogeneous species responses in moss-associated nitrogen fixation across contrasting sites, with higher rates linked to northern conditions regardless of transplantation.

This heterogeneity likely reflects shifts in epiphytic microbial consortia. Consistent with prior findings (Holland-Moritz *et al*. 2021, Stuart *et al*. 2021), we found that moss-associated microbial communities were strongly structured by host identity (Figure 4A) and that those differences in microbiomes persisted through the transplantation (Figure 4B and 4C). Transplantation did produce subtle ASV-level shifts in the relative abundance of a small subset of taxa within otherwise stable communities over the one-year study period (Figure 5C). These included a small number of canonical nitrogen-fixing cyanobacteria, notably from the genera *Nostoc* and *Stigonema* (Figure 5), which showed similar relative abundances at both home sites but shifted when transplanted, indicating conditional responses to environmental change, either directly or indirectly via host-mediated effects. Consistent with previous studies, changes in nitrogen fixation were linked to shifts in a small, environmentally sensitive subset of taxa, even though community composition remained largely unchanged (DeLuca *et al*. 2007, Allison and Martiny 2008, Gundale *et al*. 2012a). However, because we relied on 16S rRNA gene amplicon data, our inferences are limited to compositional shifts and cannot directly resolve changes in absolute abundance, functional gene context, or transcriptional activity of transplant-responsive microbes (Poretsky *et al*. 2014, Gloor *et al*. 2017). Together, these findings align with growing evidence in which microbial processes respond rapidly to environmental variation despite taxonomic composition stability (DeLuca *et al*. 2007, Stuart *et al*. 2021).

After accounting for moss species in the mixed-effects model, untransplanted Toolik mosses had an almost identical mean nitrogen-fixation rate to Healy mosses transplanted to Toolik. These observations support recent findings that colder temperatures do not necessarily suppress rates of nitrogen fixation (Andersen *et al*. 2025), and higher air and soil temperatures do not necessarily correlate with peaks in fixation activity. However, once species were analyzed separately, it became clear that not all host species fixation activities responded identically to changing conditions. Given the relatively short time interval between transplantation and measurements, it is difficult to disentangle response timing from response magnitude. The moderate response seen for *A. turgidum* may reflect a slower response rate relative to *P. schreberi*. However, large experimental manipulations of temperature (∼5.7 °C) have demonstrated that changes in nitrogen fixation associated to *P. schreberi* and *H. splendens* can occur within weeks (Gundale *et al*. 2012a). In prior work near these study sites, incubation temperature was not identified as an important predictor of nitrogen fixation (Stuart *et al*. 2021). Yet, during this particular study, incubation conditions differed between sites, with higher maximum and mean temperatures at Healy than Toolik. Importantly, temperatures at both sites did not reach levels expected to inhibit nitrogen fixation (Zielke *et al*. 2002; Gundale *et al*. 2012a; Jean *et al*. 2012).

In general, the response of high latitude moss species to variation in temperature is heterogeneous (Wahren *et al*. 2005, Hudson and Henry 2010, Prather *et al*. 2019, Alatalo *et al*. 2020). Even within species, genetic variation can lead to heterogeneity (Cronberg *et al*. 1997, Pauls *et al*. 2013), although we cannot estimate this in our experiment given the small sample sizes. Previous experimental temperature increases have observed either no change or a decrease of nitrogen-fixation rates in association with *H. splendens* (Sorensen and Michelsen 2011, Gundale *et al*. 2012a, Sorensen *et al*. 2012). Similarly, studies that quantified biomass or percent cover also saw steady or decreasing *H. splendens* cover with warming (Wahren *et al*. 2005, Alatalo *et al*. 2020), which could be related to N acquisition declines. In contrast, *P. schreberi* often responds rapidly to warming both in biomass and rates of associated nitrogen fixation (Gundale *et al*. 2012a, Deane-Coe *et al*. 2015, Rousk *et al*. 2017). Transcriptome analysis of *A. turgidum* revealed more genes related to oxidative stress from heat than cold stress as evidence of their acclimation to cold ecosystems (Liu *et al*. 2010). Biomass of *A. turgidum* has responded inconsistently to increases in air temperature (Wahren *et al*. 2005, Hudson and Henry 2010, Sorensen and Michelsen 2011), but nitrogen-fixation rates have decreased relative to the control after 20 years of experimental warming (Sorensen *et al*. 2012). In short, responses to temperature changes interact both with moss species and likely with other environmental conditions given the divergence of both biomass and nitrogen fixation rates in response to temperature alterations.

Other climatic variables with the potential to influence moss-associated nitrogen fixation include moisture conditions and light availability. The amount of precipitation was higher at Healy than Toolik at the time of our measurements, but previous studies report precipitation frequency as a more important rate variation driver than amount alone, which was similar between Toolik and Healy (Figure 7B) (Markham 2009, Gundale *et al*. 2012b). Light availability is known to affect nitrogen fixation given the prevalence of phototrophic nitrogen-fixing microbes (Basilier 1980, Gentili *et al*. 2005, Gundale *et al*. 2012a). Higher average daily PAR at Toolik, likely reflecting longer daylight hours at 68°N, may therefore contribute to higher nitrogen fixation activity at this site (Figure 7A). Relative humidity was also higher at Healy than Toolik (Figure S11) and has been shown to be as influential as temperature in regulating nitrogen fixation, with higher humidity associated with increased rates (Rousk *et al*. 2013, 2018). *P. schreberi* may also be better adapted to drier conditions than *H. splendens* due to efficient internal apoplasmic transportation of water along a central strand of hyoid cells (Sokołowska *et al*. 2017). As humidity declines, *P. schreberi* can increase long-distance internal water transportation within the moss body, a trait that may enhance moss hydration while limiting water availability to epiphytic nitrogen fixers (Sokołowska *et al*. 2017). Consequently, in drier conditions mosses could allocate water to survival while restricting available moisture for nitrogen fixation.

It remains possible that surrounding plant community composition indirectly or directly influenced observed nitrogen fixation measurements (Figure 6). Although vascular and nonvascular composition did not change within one year of transplantation, vascular plant assemblages differed between the two sites (Figure 6). Assemblages were structured by home site, with *Rubus chamaemorus* and *Rhododendron tomentosum* characterizing Healy communities and *Cassiope tetragona* dominating at Toolik. These species differ in growth form, leaf area, and phenology, suggesting that vascular plant composition may influence local light penetration and litter inputs rather than responding dynamically on the timescale of this experiment. Such structural differences are relevant to moss-associated nitrogen fixation as both light availability and litter deposition can modify microclimatic conditions and resource availability at the bryosphere scale (Zielke *et al*. 2002, Sorensen and Michelsen 2011, Gundale *et al*. 2012a, Jean *et al*. 2020). Deciduous shrubs with larger leaves, such as *R. chamaemorus*, may reduce light penetration and increase litter inputs relative to evergreen, low-stature species like *C. tetragona*, potentially contributing to site-level differences in nitrogen fixation rates. Importantly, these effects reflect persistent environmental filtering associated with site identity rather than short-term changes in plant community composition. Further, changes in moss biomass or community composition in boreal and tundra ecosystems are typically observed only after extended monitoring periods in manipulative climate experiments (Wahren *et al*. 2005, Lang *et al*. 2012, Turetsky *et al*. 2012, Deane-Coe *et al*. 2015, Alatalo *et al*. 2020), although exceptions have been reported in prior studies (Hudson and Henry 2010, Prather *et al*. 2019).

### Conclusion

Here we have shown that host identity influences moss-associated nitrogen fixation in a one-year reciprocal transplant experiment, acting both directly (through environment-driven physiological change) and indirectly (through shifts in microbial community composition). This specificity may result from differences in microbiome structure and resilience to changing conditions (Ininbergs *et al*. 2011, Jean *et al*. 2020, Holland-Moritz *et al*. 2021, Gentili *et al*. 2005), as well as differences in moss anatomy, microhabitat, and community structure that influence bryosphere temperature and moisture (Elumeeva *et al*. 2011). Given strong evidence that mosses recruit specific bacteria to acquire microbe-fixed nitrogen (Berg *et al*. 2013, Bay *et al*. 2013, Rousk *et al*. 2016, Stuart *et al*. 2020), variation in (uncharacterized) exported moss metabolites may further drive variation in fixation rates. Importantly, decreases in nitrogen supply in response to changing environmental conditions could provide a mechanism for observed declines in moss diversity under climate change (Lang *et al*. 2012, Alatalo *et al*. 2020). Changes in nitrogen inputs therefore may depend on the changing abundances of moss species and how their associated fixation rates respond to shifting conditions (Ininbergs *et al*. 2011, Jean *et al*. 2020, Holland-Moritz *et al*. 2021) supporting the need for a higher resolution monitoring of moss cover in high-latitudes ecosystems (Lett *et al*. 2022). Multi-year transplant studies that incorporate functional assays (*e.g.*, nifH assays, metagenomics, or activity-based approaches) will be critical to this effort, and to determine how changes in moss cover or biomass influence other factors like rates of new nitrogen inputs, decomposition rates, and understory heat fluxes in these areas (Blok *et al*. 2011, Turetsky *et al*. 2012).

## Data Availability

Nitrogen-fixation data are available on the Bonanza Creek LTER Data Catalog (Stuart, Julia; Mack, Michelle Cailin; Miller, Samantha. 2022). Reciprocal transplant of mosses from Arctic tundra to alpine tundra and associated nitrogen-fixation rates, Bonanza Creek LTER - University of Alaska Fairbanks. BNZ:807, http://www.lter.uaf.edu/data/data-detail/id/807). Raw 16S amplicon sequencing data is available at NCBI Sequence Read Archive under BioProject accession PRJNA1075763. Processed amplicon data, analysis code, and associated weather and plant community datasets are available at https://zenodo.org/records/19100844.

## Author Contributions

SFM, NF, and MCM obtained funding. SFM, NF, MCM, JMS, and HHM contributed to designing the research. SFM, MCM, JMS, and HHM performed field work. JMS, RSK, EL, and MH performed processing, laboratory, and statistical work. JMS, RSK, and SFM wrote the manuscript, with all other authors contributing equally to the editing of the manuscript.

## Acknowledgments

Weather data for Healy were provided by the Bonanza Creek LTER, a partnership between the University of Alaska Fairbanks and the U.S. Forest Service. Significant funding for the collection of these data was provided by the National Science Foundation Long-Term Ecological Research program (NSF Grant numbers DEB-1636476, DEB-1026415, DEB-0620579, DEB-0423442, DEB-0080609, DEB-9810217, DEB-9211769, DEB-8702629) and by the USDA Forest Service, Pacific Northwest Research Station (Agreement # RJVA-PNW-01-JV-11261952-231). Toolik weather datasets were provided by the Toolik Lake Field Station Environmental Data Center. This material is based upon work supported by the National Science Foundation under grant #1623461. Funding for experiment was provided by the National Science Foundation Division of Environmental Biology Award 1542586. Further graduate student support was provided by the ARCS Scholarship Award and Northern Arizona University. Thanks to the Bonanza Creek LTER, Arctic LTER, University of Alaska Fairbanks, and Toolik Field Station for their assistance and facilities. Thanks to Haley Dunleavy, Kyoko Okano, Julia Butler, Juan Contreras, Angela Roles, Emily Brooke, Heidi Rodenhizer, Xanthe Walker, and Henry Grover for assistance. Additional thanks to Rebecca S. Key for providing figure and moss illustrations.

