## Supplemental Figures and Tables for "Moss Transplants in the Tundra Reveal Host-Specific Microbiomes and Nitrogen Fixation Responses"

**S1:** 16S Sequencing Depth Across Samples
**S2:** Number of ASVs Across Samples
**S3:** Analysis of Variance for ASV Samples.
**S4:** Tukey HSD and Cohen's D for Diversity Metrics.
**S5:** Diversity Metrics Before and After Rarefying 16S Amplicons
**S6:** Relative Percent Abundance at Phylum-Level for Each Moss Species
**S7:** Hierarchical Clustering of ASV Samples.
**S8:** Differential Abundance Analysis of ASVs Across Treatment Groups
**S9:** Species Associations with Non-Vascular Plant Community Composition
**S10:** Significant Species Associations with Vascular Plant Community
**S11:** Average Daily Relative Humidity Leading Up to Nitrogen Fixation Measurements

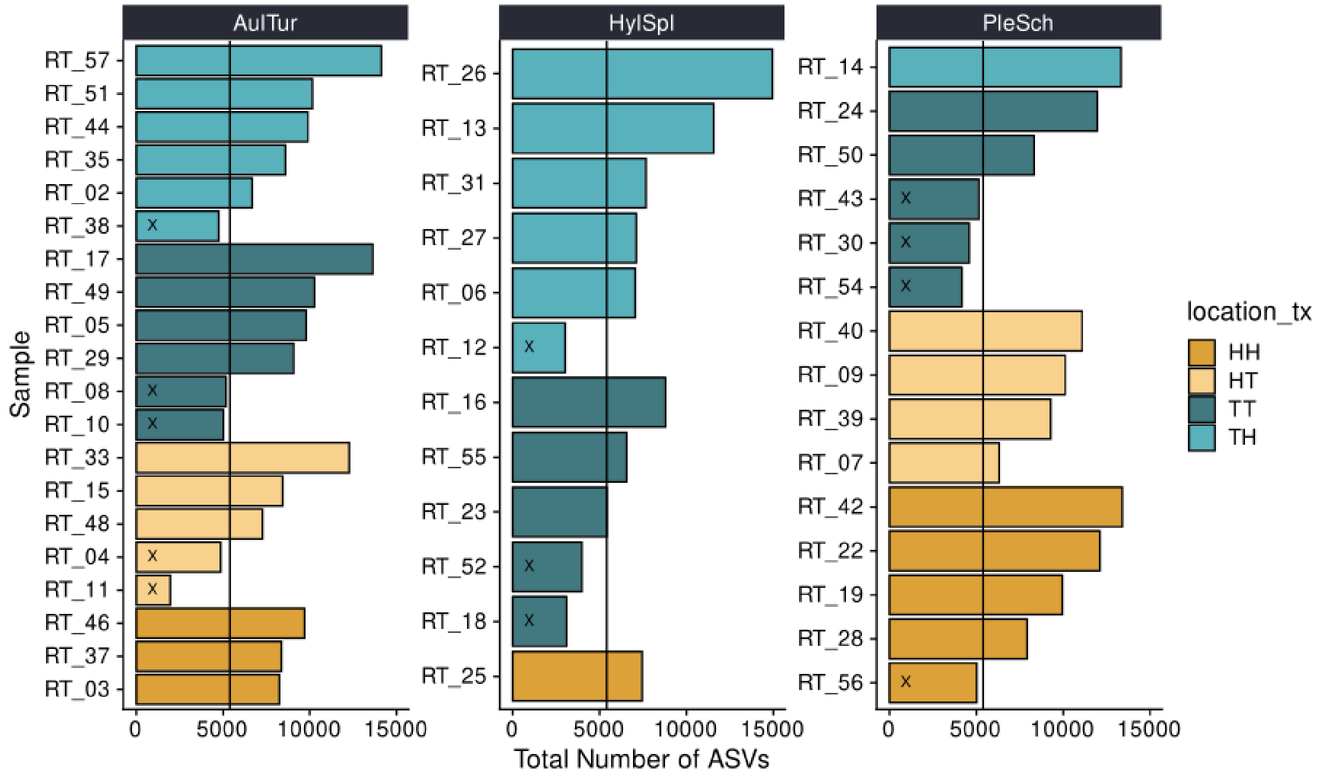


**S1.** 16S Sequencing Depth Across Samples. This bar chart shows the number of amplicon sequence variants (ASVs) per sample for three moss species: *Aulcomium turgidum* (AulTur), *Hylocomium splendens* (HylSpl), and *Pleurozium schreberi* (PleSch). The y-axis represents samples, and the x-axis represents ASV counts. The black line indicates the rarefy threshold, which was set at 5400 reads. Samples below this threshold (marked with 'X') were excluded from the 16S analysis. Bars are color-coded by location treatment: HH = Healy Home, HT = Healy Away, TT = Toolik Home, and TH = Toolik Away. Each moss species contained at least three replicates per treatment group, except for *P. schreberi*'s Toolik Home (n = 2) and Toolik Away (n = 1) treatments, as well as *H. splendens*'s Healy Home treatment (n = 1) and Healy Away (n = 0).

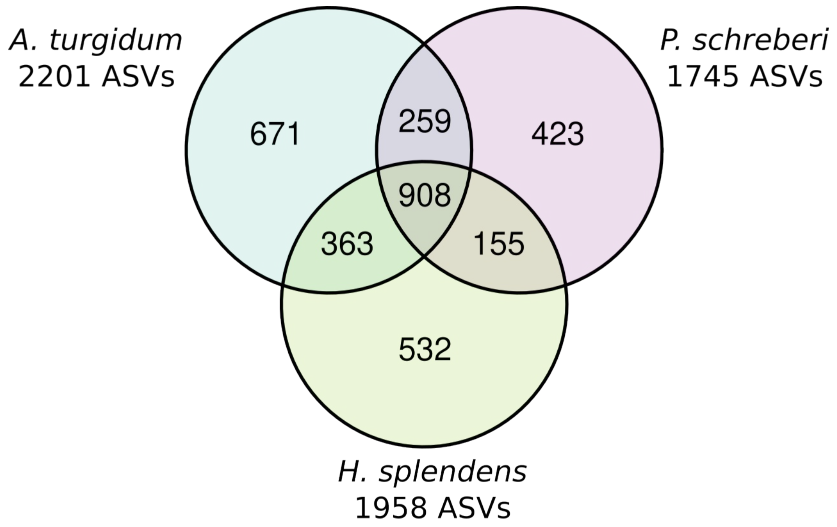


**S2.** Number of ASVs Across Samples. A Venn diagram showing the total and shared ASVs among Aulacomnium turgidum (2201), Pleurozium schreberi (1745), and Hylocomium splendens (1958). The rarefaction threshold was set to 5400 reads.

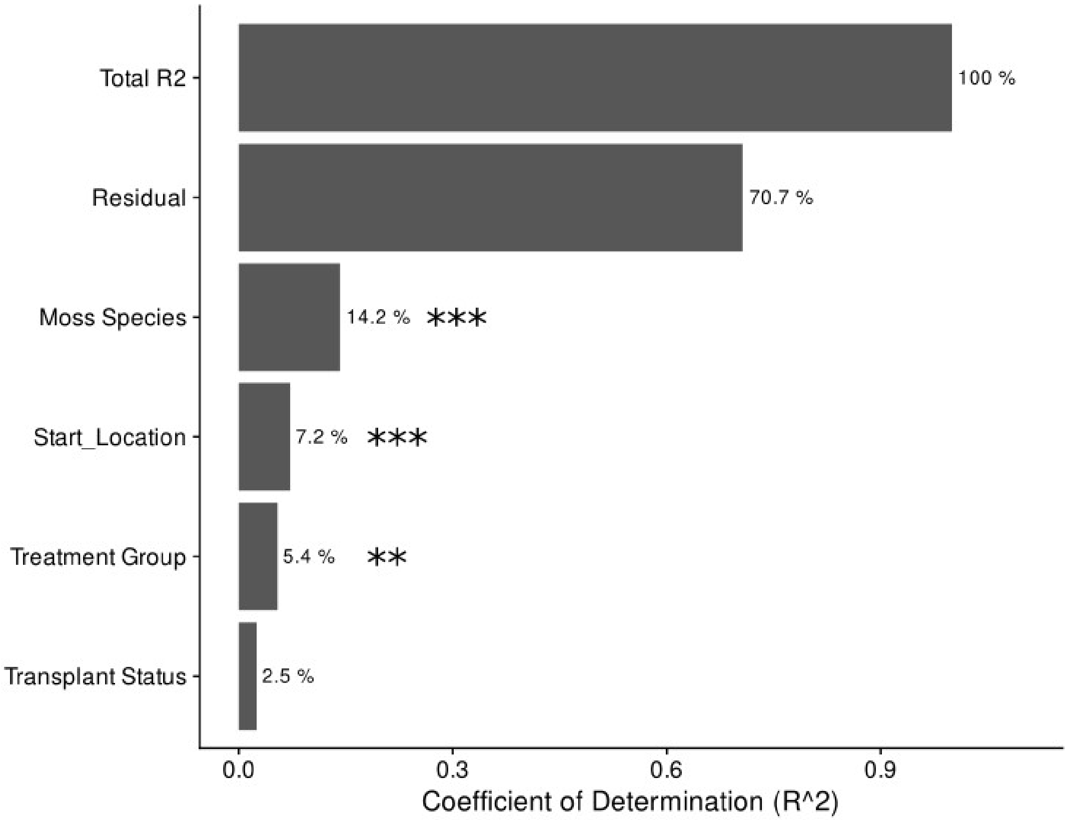


**S3**. Analysis of Variance for ASV Samples. Bar plot showing $R^2$ values from Permutational Multivariate Analysis of Variance (PERMANOVA) based on Bray-Curtis dissimilarity. The effect of Moss Species (*A. turgidum, H. splendens*, and *P. schreberi*), Start Location, treatment group (Heal Home, Healy Away, Toolik Home, Toolik Away), transplant status (transplanted and untransplanted), and residual (ie. unexplained variance). The x-axis contains the $R^2$ values, indicating the proportion of variance explained by each factor. These proportions are in percentage and are displayed at the end of the corresponding bars. 'Total $R^2$' accounts for all cumulative variance from the categories below it, and asterisks denote level of significance with p-values < 0.05.

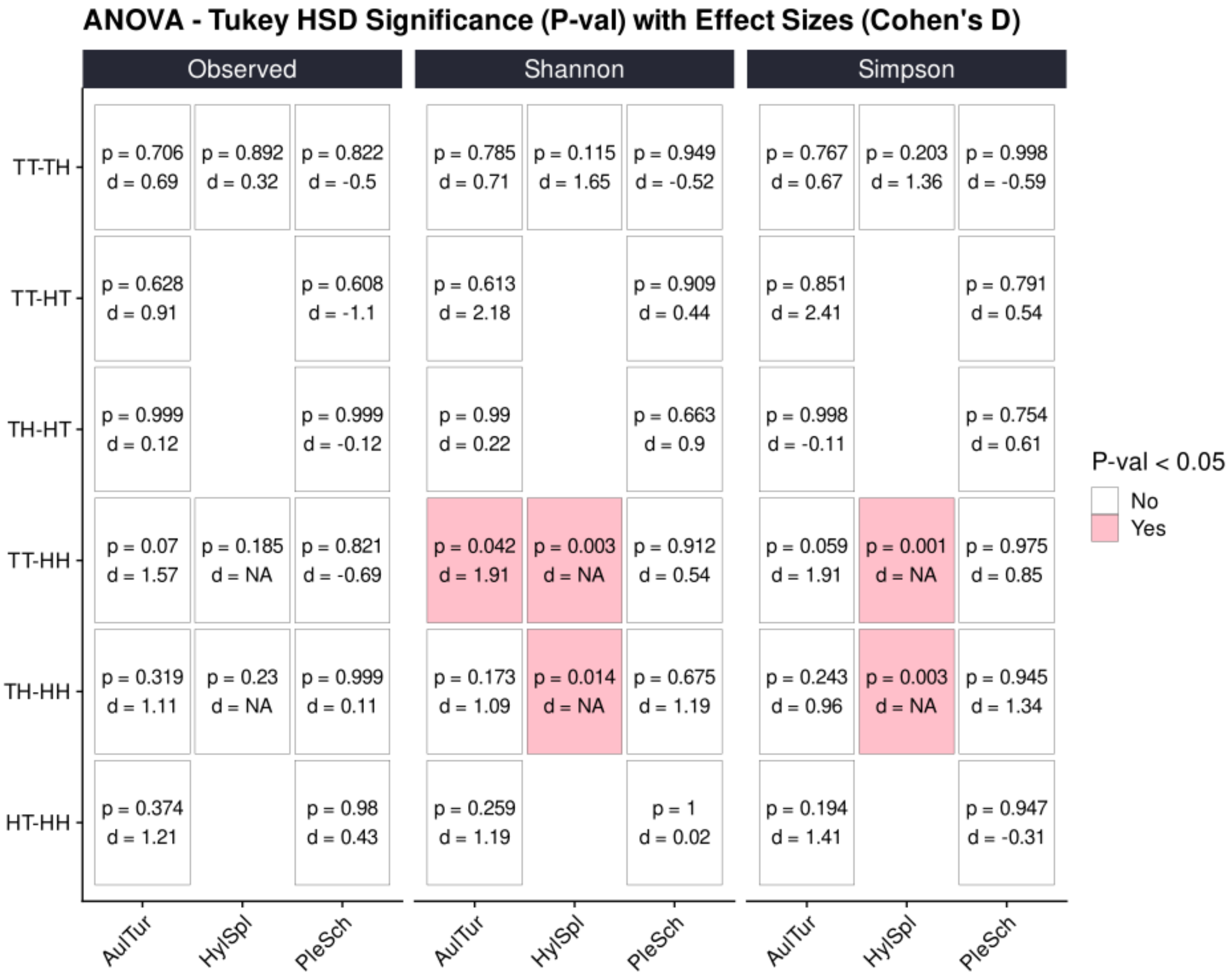


**S4.** Tukey HSD and Cohen's D for Diversity Metrics. A heatmap showing ANOVA followed by Tukey's Honest Significant Difference (HSD) tests for three diversity measures: Observed ASVs, Shannon diversity, and Simpson index, across samples from three moss species: *Aulacomnium turgidum* (AuTur), *Hylocomium splendens* (HylSpl), and *Pleurozium schreberi* (PleSch). The comparisons are made between 'home' treatments (TT = Toolik-Toolik and HH = Healy-Healy) and 'away' treatments (TH = Toolik-Healy and HT = Healy-Toolik). The p-values indicate the significance of the differences between treatments, with values less than 0.05 highlighted in pink. Effect sizes are presented as Cohen's D values. In instances where a Cohen's D value could not be calculated due to < 2 samples in one treatment group, an NA is present.  *H. splendens* only contained 1 sample in the 'HH' group.

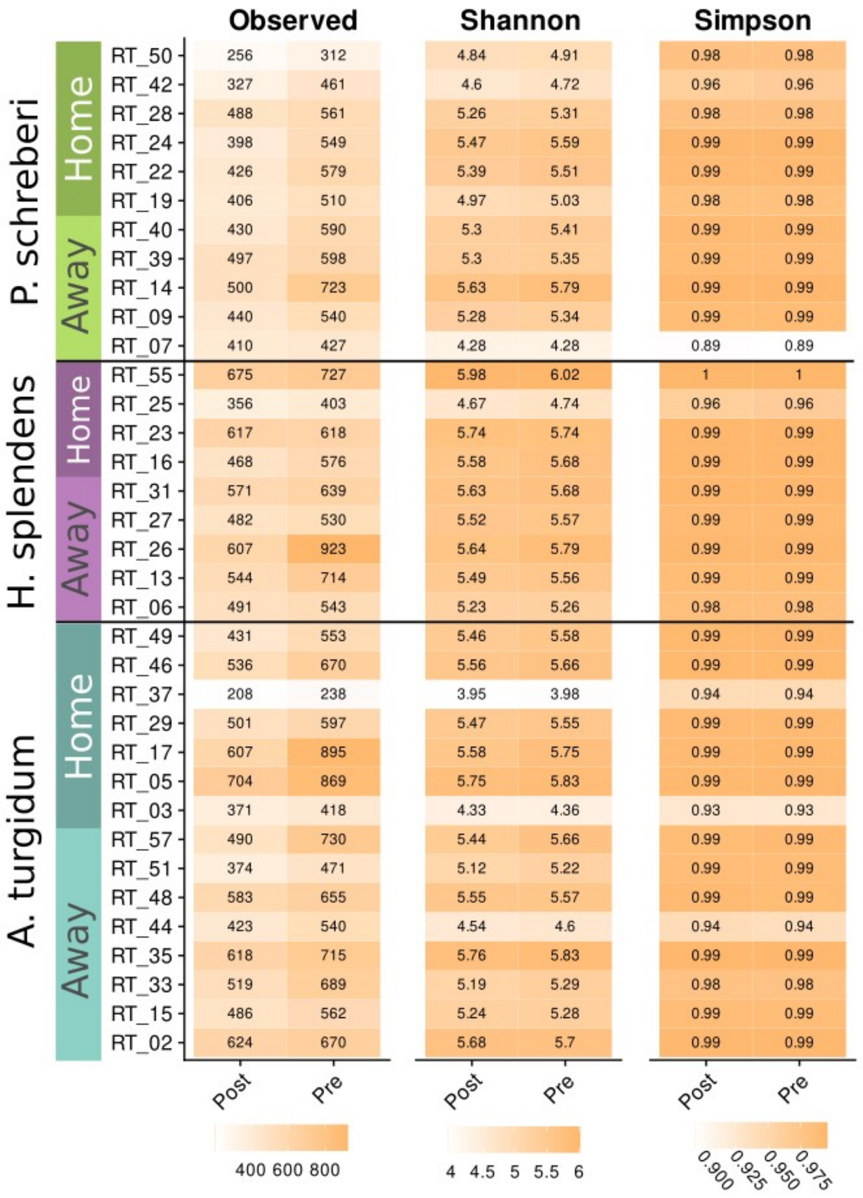


**S5.** Diversity Metrics Before and After Rarefying 16S Amplicons. The following heatmap illustrates the diversity measures – Observed ASVs, Shannon diversity, and Inverse Simpson index – across samples from three moss species. A. turgidum samples are colored teal, H. splendens in purple, and P. schreberi in green. The samples are categorized into 'home' treatments (contains TT = Toolik-Toolik and HH = Healy-Healy samples) and 'away' treatments (contains TH = Toolik-Healy and HT = Healy-Toolik samples). The diversity values of samples in the y-axis are presented in two columns: before rarefying (Pre) and after rarefying (Post). The color scale is based on the range of diversity measures, with higher values indicating greater diversity and colored yellow. Notably, post-rarefied sample values are generally lower but show almost similar values as the pre-rarefied samples. The slightly lower values reflect the effect of standardized read depth on diversity estimation and how normalization of sequencing depth has minimal effect on this particular dataset.

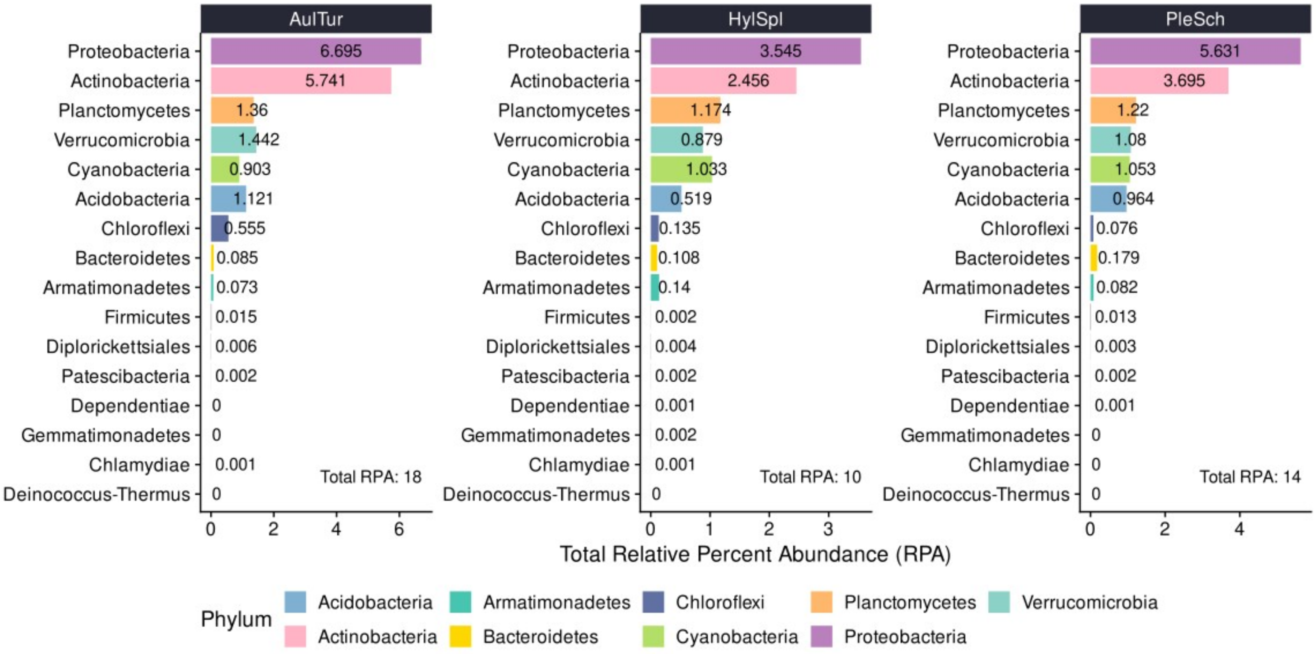


**S6.** Relative Percent Abundance at Phylum-Level for Each Moss Species. Bar charts shows the total relative percent abundance of microbial phyla associated with three moss species: *Aulcomium turgidum* (AulTur), *Hylocomium splendens* (HylSpl), and *Pleurozium schreberi* (PleSch). ASVs classified as ‘Chloroplast’ or ‘Rickettsiales’ were then treated as contaminants and removed from the 16S microbial analysis. The x-axis shows relative percent abundance values, with relative percent abundance values printed to the right of each bar in the plot. Values less than 0.001 are represented as zero due to rounding. The total relative percent abundance, summed across all phyla, are located to the bottom right of each plot.

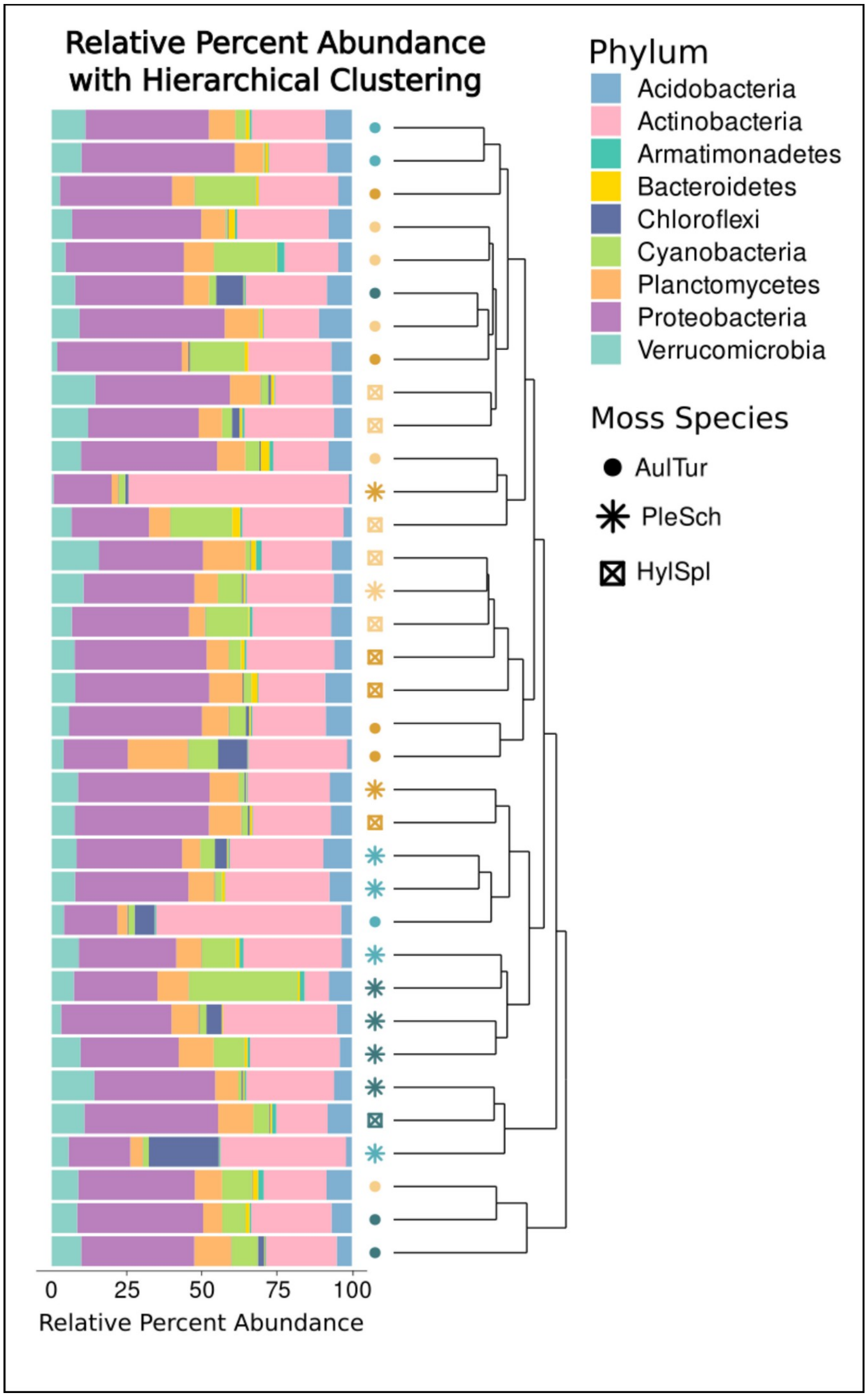


**S7.** Hierarchical Clustering of ASV Samples. Each horizontal bar represents an individual sample, with colors indicating 16S phylums. The clustering dendrogram is to the right of sample bars, clustering samples based on microbial composition similarity at phylum-level. The dendrogram tree tips are shaped according to moss species: *Aulacomnium turgidum* (AulTur; circles), *Pleurozium schreberi* (PleSch; asterisks), and *Hylocomium splendens* (HylSpl; squares with Xs inside). Shaped tip points are also color-coded based on treatment group: Toolik Home (TT; dark teal), Toolik Away (TH; light teal), Healy Home (HH; dark gold), and Healy Away (HT; light gold).

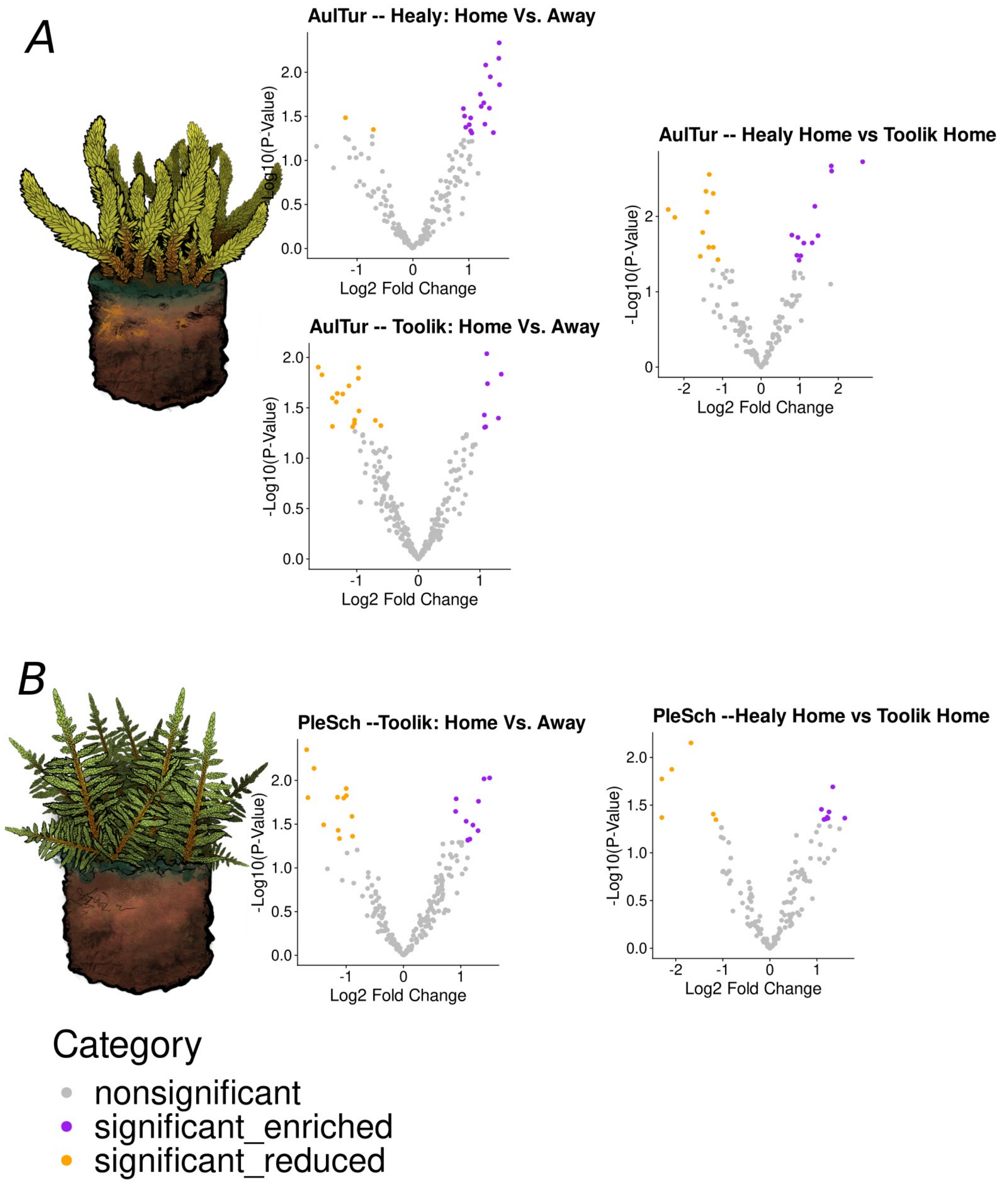


**S8.** Differential Abundance Analysis of ASVs Across Treatment Groups. **(A)** *Aulacomnium turgidum* and **(B)** *Pleurozium schreberi* volcano plots from ANCOM-BC analysis comparing ASVs between home controls and reciprocal transplant treatments. Each plot displays log2 fold change on the x-axis and -log10(p-value) on the y-axis. ASVs significantly enriched (upregulated) in transplant conditions are highlighted in purple, while significantly reduced (downregulated) ASVs are shown in orange. Nonsignificant ASVs are represented in grey.

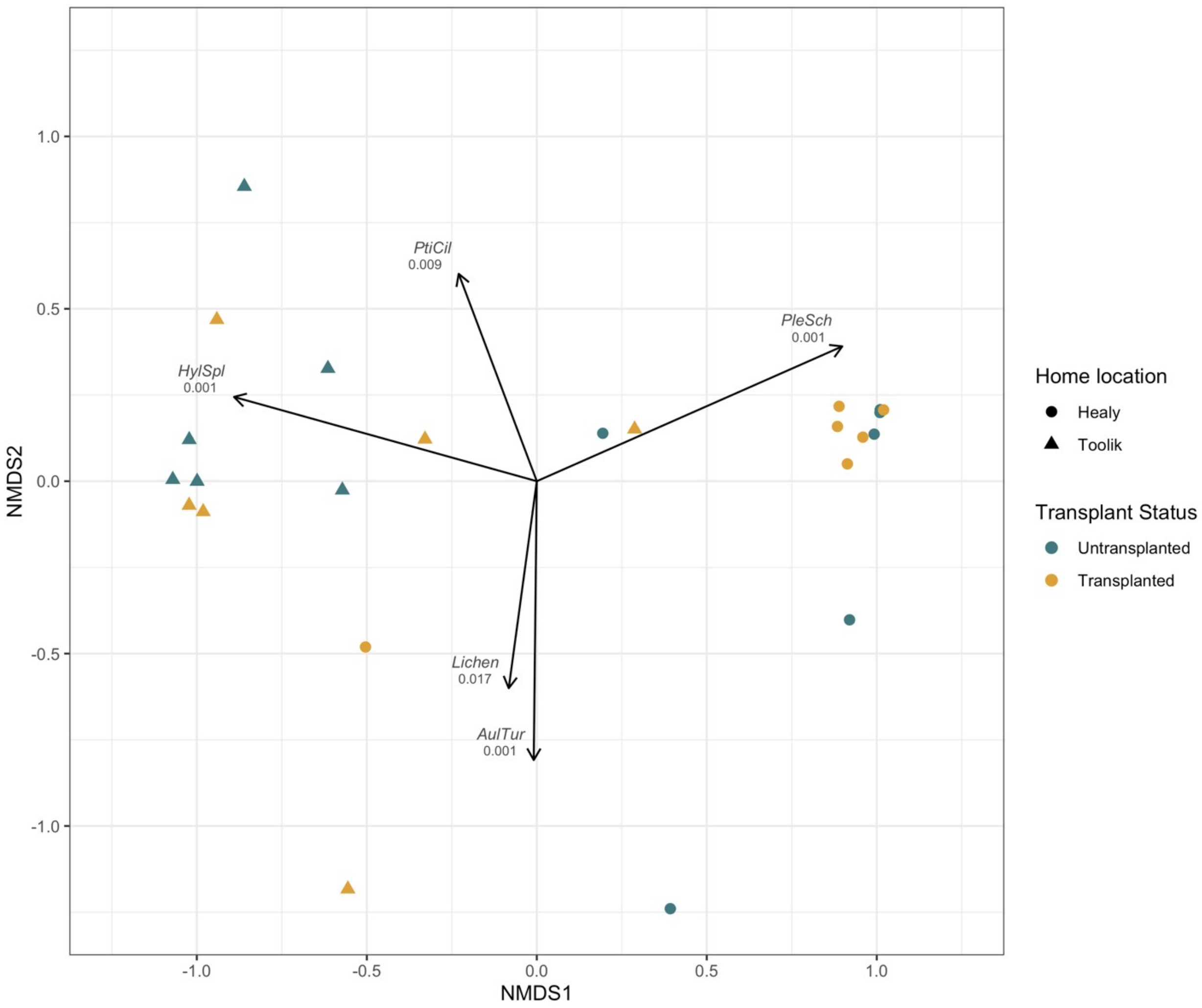


**S9.** Species Associations with Non-Vascular Plant Community Composition. Vectors of species fit on non-vascular plant community composition from both experimental sites. Vectors are pictured if the P value was less than 0.05, and the P values are included next to the species code in the graph. Species codes are the first three letters of the genus followed by the first three letters of the species. If species were not differentiated within a genus, *Spp* appears in place of the species code.

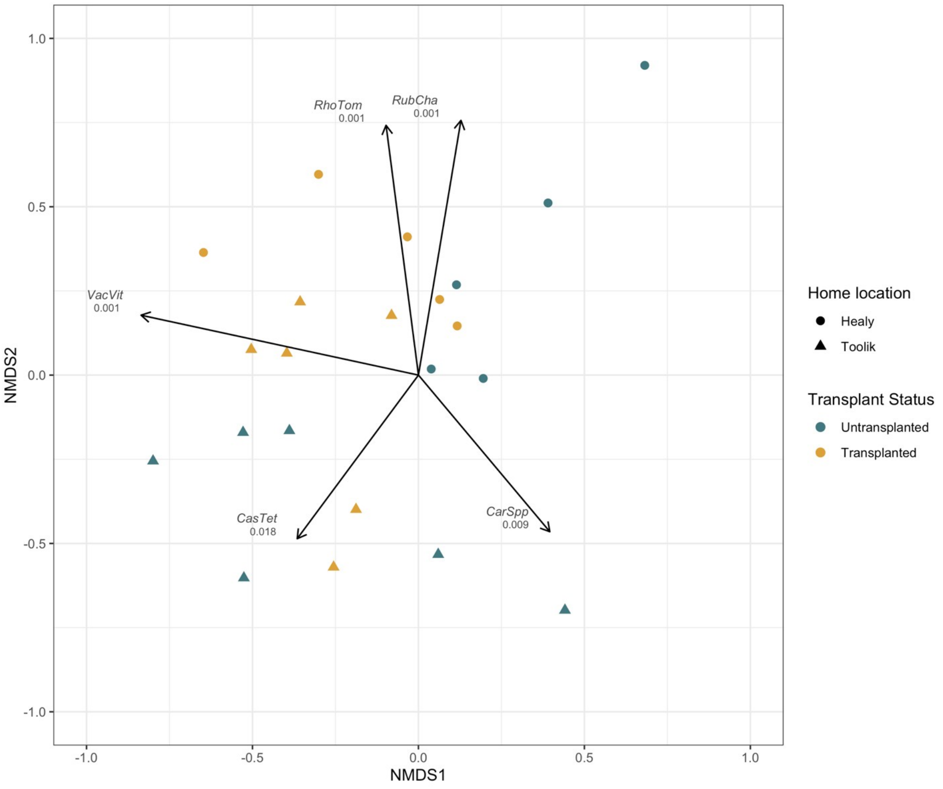


**S10.** Significant Species Associations with Vascular Plant Community. Vectors of species fit on vascular plant community composition from both experimental sites. Vectors are pictured if the P value was less than 0.05, and the P values are included next to the species code in the graph. Species codes are the first three letters of the genus followed by the first three letters of the species. If species were not differentiated within a genus, *Spp* appears in place of the species code.

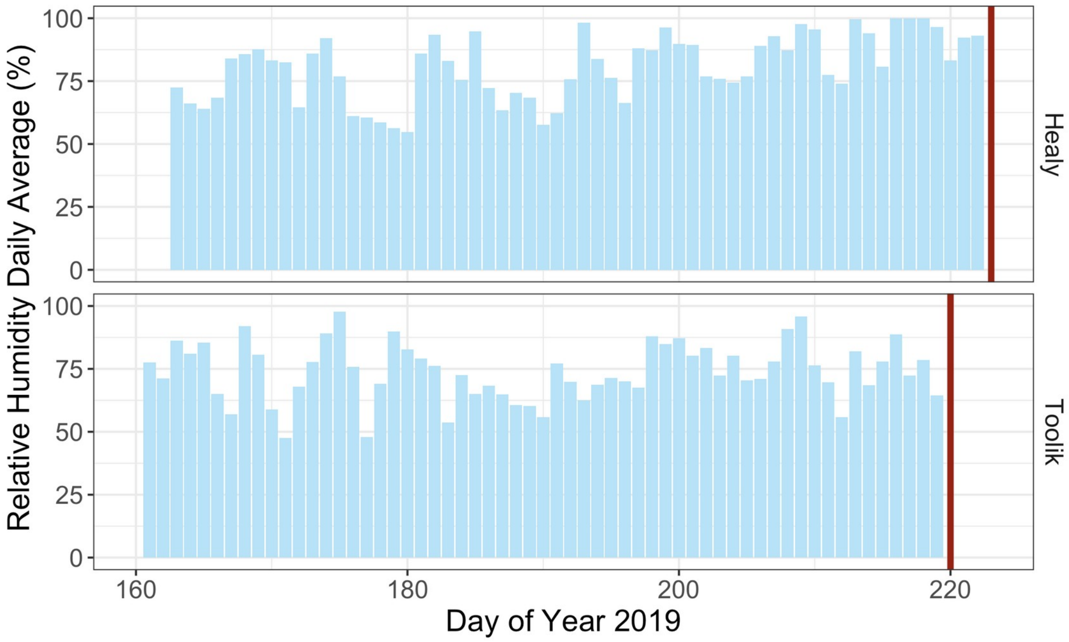


**S11**. Average Daily Relative Humidity Leading Up to Nitrogen Fixation Measurements. Average daily relative humidity in the growing season leading up to nitrogn fixation measurements. Dates of incubation are indicated by a red bar on each graph. The *x*-axis is the Julian day of year.

**Supplemental Tables**

**S1:** Significant Differentially Abundant ASVs for *Aulacomnium turgidum*

**S2:** Significant Differentially Abundant ASVs for *Pleurozium schreberi*

**S1.** Significant Differentially Abundant ASVs for *Aulacomnium turgidum*. Significant Differentially Abundant ASVs for *Aulacomnium turgidum*. This table provides the taxonomic classifications for significant ASVs corresponding to P. schreberi and identified through Analysis of Composition of Microbiomes with Bias Correction (ANCOM-BC). Each row represents an ASV, with columns detailing log2 fold change, adjusted p-value (BH corrected), ANCOMBC comparison group, and regulation status (enriched or reduced). Taxonomy classifications are provided from Class to Genus, with unknown classifications marked as "NA."

### *A. turgidum* ANCOM-BC

| ASV ID | Taxonomic Classification | Log2 Fold Change | Adjusted P-Value | Condition | Regulation |
|---|---|---|---|---|---|
| ASV_26 | Proteobacteria; Gammaproteobacteria; WD260; NA; NA | 0.9834712 | 0.038133410 | Controls | enriched |
| ASV_57 | Planctomycetes; Planctomycetacia; Isosphaerales; Isosphaeraceae; Singulisphaera | 1.8196215 | 0.002137429 | Controls | enriched |
| ASV_58 | Proteobacteria; Alphaproteobacteria; Acetobacterales; Acetobacteraceae; NA | 0.7976317 | 0.017708477 | Controls | enriched |
| ASV_93 | Planctomycetes; Planctomycetacia; Isosphaerales; Isosphaeraceae; Singulisphaera | 1.8279005 | 0.002495028 | Controls | enriched |
| ASV_330 | Proteobacteria; Alphaproteobacteria; Rhizobiales; Beijerinckiaceae; Roseiarcus | 0.9551256 | 0.018990271 | Controls | enriched |
| ASV_355 | Proteobacteria; Alphaproteobacteria; Acetobacterales; Acetobacteraceae; NA | 1.0274830 | 0.033293683 | Controls | enriched |
| ASV_379 | Verrucomicrobia; Verrucomicrobiae; Chthoniobacterales; Xiphinematobacteraceae; Candidatus_Xiphinematobacter | 1.4757776 | 0.017998712 | Controls | enriched |
| ASV_526 | Proteobacteria; Alphaproteobacteria; Acetobacterales; Acetobacteraceae; NA | 1.3239007 | 0.022432461 | Controls | enriched |
| ASV_835 | Verrucomicrobia; Verrucomicrobiae; Chthoniobacterales; Xiphinematobacteraceae; Candidatus_Xiphinematobacter | 2.6321793 | 0.001879263 | Controls | enriched |
| ASV_969 | Acidobacteria; Acidobacteriia; Acidobacteriales; Acidobacteriaceae_(Subgroup_1); Acidipila | 1.3937356 | 0.007337569 | Controls | enriched |
| ASV_1574 | Proteobacteria; Alphaproteobacteria; Acetobacterales; Acetobacteraceae; NA | 1.1073392 | 0.022583165 | Controls | enriched |
| ASV_1814 | Proteobacteria; Alphaproteobacteria; Acetobacterales; Acetobacteraceae; NA | 0.9274694 | 0.032850175 | Controls | enriched |
| ASV_35 | Actinobacteria; Actinobacteria; Frankiales; Frankiaceae; Jatrophihabitans | -1.3541100 | 0.025629678 | Controls | reduced |
| ASV_92 | Actinobacteria; Thermoleophilia; Solirubrobacterales; Solirubrobacteraceae; Conexibacter | -1.2391589 | 0.025664868 | Controls | reduced |
| ASV_356 | Actinobacteria; Acidimicrobiia; IMCC26256; NA; NA | -1.1168556 | 0.037520616 | Controls | reduced |
| ASV_537 | Actinobacteria; Acidimicrobiia; IMCC26256; NA; NA | -1.5756193 | 0.033835425 | Controls | reduced |
| ASV_816 | Actinobacteria; Actinobacteria; Frankiales; Acidothermaceae; Acidothermus | -2.2353189 | 0.010291542 | Controls | reduced |
| ASV_938 | Actinobacteria; Thermoleophilia; Solirubrobacterales; Solirubrobacteraceae; Conexibacter | -1.4265511 | 0.004640292 | Controls | reduced |
| ASV_1146 | Actinobacteria; Thermoleophilia; Solirubrobacterales; Solirubrobacteraceae; Conexibacter | -1.3422367 | 0.002771610 | Controls | reduced |
| ASV_1170 | Actinobacteria; Actinobacteria; Streptomycetales; Streptomycetaceae; Streptacidiphilus | -1.3972788 | 0.008755898 | Controls | reduced |
| ASV_1453 | Actinobacteria; Thermoleophilia; Solirubrobacterales; Solirubrobacteraceae; Conexibacter | -2.4059768 | 0.008087352 | Controls | reduced |
| ASV_1486 | Actinobacteria; Acidimicrobiia; IMCC26256; NA; NA | -1.5119940 | 0.016312594 | Controls | reduced |
| ASV_3582 | Actinobacteria; Acidimicrobiia; IMCC26256; NA; NA | -1.2395221 | 0.004956968 | Controls | reduced |
| ASV_39 | Verrucomicrobia; Verrucomicrobiae; Chthoniobacterales; Chthoniobacteraceae; Chthoniobacter | 1.5380013 | 0.006940970 | HealyRT | enriched |
| ASV_57 | Planctomycetes; Planctomycetacia; Isosphaerales; Isosphaeraceae; Singulisphaera | 1.5506690 | 0.013822260 | HealyRT | enriched |
| ASV_80 | Proteobacteria; Alphaproteobacteria; Caulobacterales; Caulobacteraceae; NA | 0.9024309 | 0.025810930 | HealyRT | enriched |
| ASV_83 | Acidobacteria; Acidobacteriia; Acidobacteriales; Acidobacteriaceae_(Subgroup_1); Granulicella | 0.9241255 | 0.031406151 | HealyRT | enriched |
| ASV_93 | Planctomycetes; Planctomycetacia; Isosphaerales; Isosphaeraceae; Singulisphaera | 1.5435529 | 0.004625560 | HealyRT | enriched |
| ASV_179 | Proteobacteria; Gammaproteobacteria; WD260; NA; NA | 1.4399588 | 0.048355403 | HealyRT | enriched |
| ASV_255 | Proteobacteria; Alphaproteobacteria; Rhizobiales; Beijerinckiaceae; Roseiarcus | 1.2073286 | 0.017707401 | HealyRT | enriched |
| ASV_263 | Proteobacteria; Alphaproteobacteria; Rhizobiales; Beijerinckiaceae; 1174-901-12 | 1.0333745 | 0.032932678 | HealyRT | enriched |
| ASV_305 | Acidobacteria; Acidobacteriia; Acidobacteriales; Acidobacteriaceae_(Subgroup_1); NA | 1.3848003 | 0.011238640 | HealyRT | enriched |
| ASV_355 | Proteobacteria; Alphaproteobacteria; Acetobacterales; Acetobacteraceae; NA | 1.3686657 | 0.025495401 | HealyRT | enriched |
| ASV_461 | Actinobacteria; Actinobacteria; Corynebacteriales; Mycobacteriaceae; Mycobacterium | 1.2888370 | 0.038761786 | HealyRT | enriched |
| ASV_526 | Proteobacteria; Alphaproteobacteria; Acetobacterales; Acetobacteraceae; NA | 1.0085797 | 0.039261192 | HealyRT | enriched |
| ASV_817 | Acidobacteria; Acidobacteriia; Solibacterales; Solibacteraceae_(Subgroup_3); Candidatus_Solibacter | 1.2684845 | 0.022203329 | HealyRT | enriched |
| ASV_835 | Verrucomicrobia; Verrucomicrobiae; Chthoniobacterales; Xiphinematobacteraceae; Candidatus_Xiphinematobacter | 0.9459919 | 0.041967306 | HealyRT | enriched |
| ASV_844 | Acidobacteria; Acidobacteriia; Acidobacteriales; Acidobacteriaceae_(Subgroup_1); Granulicella | 1.2222566 | 0.024357110 | HealyRT | enriched |
| ASV_969 | Acidobacteria; Acidobacteriia; Acidobacteriales; Acidobacteriaceae_(Subgroup_1); Acidipila | 1.3033496 | 0.008266846 | HealyRT | enriched |
| ASV_1244 | Proteobacteria; Alphaproteobacteria; Acetobacterales; Acetobacteraceae; Endobacter | 1.0541600 | 0.048841532 | HealyRT | enriched |
| ASV_1687 | Proteobacteria; Alphaproteobacteria; Acetobacterales; Acetobacteraceae; Acidiphilium | 1.0342229 | 0.046043558 | HealyRT | enriched |
| ASV_1146 | Actinobacteria; Thermoleophilia; Solirubrobacterales; Solirubrobacteraceae; Conexibacter | -0.7080538 | 0.044588210 | HealyRT | reduced |
| ASV_4076 | Actinobacteria; Actinobacteria; Frankiales; Acidothermaceae; Acidothermus | -1.2067919 | 0.032750231 | HealyRT | reduced |
| ASV_7 | Proteobacteria; Gammaproteobacteria; Betaproteobacteriales; Burkholderiaceae; NA | 1.1115554 | 0.009172704 | ToolikRT | enriched |
| ASV_230 | Verrucomicrobia; Verrucomicrobiae; Chthoniobacterales; NA; NA | 1.3467284 | 0.014646212 | ToolikRT | enriched |
| ASV_507 | Actinobacteria; Actinobacteria; Frankiales; Frankiaceae; Jatrophihabitans | 1.0770599 | 0.049334906 | ToolikRT | enriched |
| ASV_651 | Actinobacteria; Actinobacteria; Frankiales; Acidothermaceae; Acidothermus | 1.0938334 | 0.048729877 | ToolikRT | enriched |
| ASV_697 | Actinobacteria; Actinobacteria; Frankiales; Nakamurellaceae; Nakamurella | 1.0722072 | 0.037277545 | ToolikRT | enriched |
| ASV_835 | Verrucomicrobia; Verrucomicrobiae; Chthoniobacterales; Xiphinematobacteraceae; Candidatus_Xiphinematobacter | 1.3007395 | 0.039996424 | ToolikRT | enriched |
| ASV_3264 | Proteobacteria; Alphaproteobacteria; Acetobacterales; Acetobacteraceae; NA | 1.1233858 | 0.018195458 | ToolikRT | enriched |
| ASV_15 | Cyanobacteria; Oxyphotobacteria; Nostocales; Nostocaceae; Nostoc_PCC-73102 | -1.0662051 | 0.048715807 | ToolikRT | reduced |
| ASV_35 | Actinobacteria; Actinobacteria; Frankiales; Frankiaceae; Jatrophihabitans | -1.3940448 | 0.025290586 | ToolikRT | reduced |
| ASV_259 | Proteobacteria; Alphaproteobacteria; Acetobacterales; Acetobacteraceae; NA | -1.3940910 | 0.048330657 | ToolikRT | reduced |
| ASV_356 | Actinobacteria; Acidimicrobiia; IMCC26256; NA; NA | -1.3155530 | 0.022751821 | ToolikRT | reduced |
| ASV_491 | Proteobacteria; Alphaproteobacteria; Acetobacterales; Acetobacteraceae; NA | -1.1264059 | 0.019110072 | ToolikRT | reduced |
| ASV_503 | Cyanobacteria; Oxyphotobacteria; Nostocales; Nostocaceae; Stigonema_SAG_48.90 | -1.0347869 | 0.041731800 | ToolikRT | reduced |
| ASV_611 | Acidobacteria; Acidobacteriia; Acidobacteriales; Acidobacteriaceae_(Subgroup_1); Granulicella | -1.0369582 | 0.045096296 | ToolikRT | reduced |
| ASV_618 | Proteobacteria; Alphaproteobacteria; Acetobacterales; Acetobacteraceae; Acidicaldus | -0.9687485 | 0.012598828 | ToolikRT | reduced |
| ASV_655 | Proteobacteria; Alphaproteobacteria; Acetobacterales; Acetobacteraceae; Acidisphaera | -0.9628915 | 0.033895510 | ToolikRT | reduced |
| ASV_731 | Actinobacteria; Acidimicrobiia; IMCC26256; NA; NA | -0.9734701 | 0.016082671 | ToolikRT | reduced |
| ASV_816 | Actinobacteria; Actinobacteria; Frankiales; Acidothermaceae; Acidothermus | -1.5637889 | 0.014853627 | ToolikRT | reduced |
| ASV_899 | Actinobacteria; Thermoleophilia; Solirubrobacterales; Solirubrobacteraceae; Conexibacter | -0.6079572 | 0.047457240 | ToolikRT | reduced |
| ASV_1066 | Cyanobacteria; Oxyphotobacteria; Nostocales; Nostocaceae; NA | -1.3323704 | 0.027649823 | ToolikRT | reduced |
| ASV_1453 | Actinobacteria; Thermoleophilia; Solirubrobacterales; Solirubrobacteraceae; Conexibacter | -0.6968134 | 0.042125565 | ToolikRT | reduced |
| ASV_1894 | Actinobacteria; Thermoleophilia; Solirubrobacterales; Solirubrobacteraceae; Conexibacter | -1.2263517 | 0.023042011 | ToolikRT | reduced |
| ASV_2598 | Actinobacteria; Actinobacteria; Frankiales; Acidothermaceae; Acidothermus | -1.6241146 | 0.012482338 | ToolikRT | reduced |

**S2.** Significant Differentially Abundant ASVs for *Pleurozium schreberi*. This table provides the taxonomic classifications for significant ASVs corresponding to P. schreberi and identified through Analysis of Composition of Microbiomes with Bias Correction (ANCOM-BC). Each row represents an ASV, with columns detailing log2 fold change, adjusted p-value (BH corrected), ANCOMBC comparison group, and regulation status (enriched or reduced). Taxonomy classifications are provided from Class to Genus, with unknown classifications marked as "NA."

### *P. schreberi* ANCOM-BC

| ASV ID | Taxonomic Classification | Log2 Fold Change | Adjusted P-Value | Condition | Regulation |
|---|---|---|---|---|---|
| ASV_26 | Proteobacteria; Gammaproteobacteria; WD260; NA; NA | 0.9834712 | 0.038133410 | Controls | enriched |
| ASV_57 | Planctomycetes; Planctomycetacia; Isosphaerales; Isosphaeraceae; Singulisphaera | 1.8196215 | 0.002137429 | Controls | enriched |
| ASV_58 | Proteobacteria; Alphaproteobacteria; Acetobacterales; Acetobacteraceae; NA | 0.7976317 | 0.017708477 | Controls | enriched |
| ASV_93 | Planctomycetes; Planctomycetacia; Isosphaerales; Isosphaeraceae; Singulisphaera | 1.8279005 | 0.002495028 | Controls | enriched |
| ASV_330 | Proteobacteria; Alphaproteobacteria; Rhizobiales; Beijerinckiaceae; Roseiarcus | 0.9551256 | 0.018990271 | Controls | enriched |
| ASV_355 | Proteobacteria; Alphaproteobacteria; Acetobacterales; Acetobacteraceae; NA | 1.0274830 | 0.033293683 | Controls | enriched |
| ASV_379 | Verrucomicrobia; Verrucomicrobiae; Chthoniobacterales; Xiphinematobacteraceae; Candidatus_Xiphinematobacter | 1.4757776 | 0.017998712 | Controls | enriched |
| ASV_526 | Proteobacteria; Alphaproteobacteria; Acetobacterales; Acetobacteraceae; NA | 1.3239007 | 0.022432461 | Controls | enriched |
| ASV_835 | Verrucomicrobia; Verrucomicrobiae; Chthoniobacterales; Xiphinematobacteraceae; Candidatus_Xiphinematobacter | 2.6321793 | 0.001879263 | Controls | enriched |
| ASV_969 | Acidobacteria; Acidobacteriia; Acidobacteriales; Acidobacteriaceae_(Subgroup_1); Acidipila | 1.3937356 | 0.007337569 | Controls | enriched |
| ASV_1574 | Proteobacteria; Alphaproteobacteria; Acetobacterales; Acetobacteraceae; NA | 1.1073392 | 0.022583165 | Controls | enriched |
| ASV_1814 | Proteobacteria; Alphaproteobacteria; Acetobacterales; Acetobacteraceae; NA | 0.9274694 | 0.032850175 | Controls | enriched |
| ASV_35 | Actinobacteria; Actinobacteria; Frankiales; Frankiaceae; Jatrophihabitans | -1.3541100 | 0.025629678 | Controls | reduced |
| ASV_92 | Actinobacteria; Thermoleophilia; Solirubrobacterales; Solirubrobacteraceae; Conexibacter | -1.2391589 | 0.025664868 | Controls | reduced |
| ASV_356 | Actinobacteria; Acidimicrobiia; IMCC26256; NA; NA | -1.1168556 | 0.037520616 | Controls | reduced |
| ASV_537 | Actinobacteria; Acidimicrobiia; IMCC26256; NA; NA | -1.5756193 | 0.033835425 | Controls | reduced |
| ASV_816 | Actinobacteria; Actinobacteria; Frankiales; Acidothermaceae; Acidothermus | -2.2353189 | 0.010291542 | Controls | reduced |
| ASV_938 | Actinobacteria; Thermoleophilia; Solirubrobacterales; Solirubrobacteraceae; Conexibacter | -1.4265511 | 0.004640292 | Controls | reduced |
| ASV_1146 | Actinobacteria; Thermoleophilia; Solirubrobacterales; Solirubrobacteraceae; Conexibacter | -1.3422367 | 0.002771610 | Controls | reduced |
| ASV_1170 | Actinobacteria; Actinobacteria; Streptomycetales; Streptomycetaceae; Streptacidiphilus | -1.3972788 | 0.008755898 | Controls | reduced |
| ASV_1453 | Actinobacteria; Thermoleophilia; Solirubrobacterales; Solirubrobacteraceae; Conexibacter | -2.4059768 | 0.008087352 | Controls | reduced |
| ASV_1486 | Actinobacteria; Acidimicrobiia; IMCC26256; NA; NA | -1.5119940 | 0.016312594 | Controls | reduced |
| ASV_3582 | Actinobacteria; Acidimicrobiia; IMCC26256; NA; NA | -1.2395221 | 0.004956968 | Controls | reduced |
| ASV_9 | Proteobacteria; Gammaproteobacteria; WD260; NA; NA | 0.9108165 | 0.022609951 | HealyRT | enriched |
| ASV_10 | Planctomycetes; Planctomycetacia; Isosphaerales; Isosphaeraceae; Singulisphaera | 1.5041065 | 0.009357755 | HealyRT | enriched |
| ASV_107 | Proteobacteria; Gammaproteobacteria; Betaproteobacteriales; Burkholderiaceae; Rhizobacter | 1.2128375 | 0.032401706 | HealyRT | enriched |
| ASV_130 | Actinobacteria; Actinobacteria; Micrococcales; Microbacteriaceae; Amnibacterium | 1.1238944 | 0.048252874 | HealyRT | enriched |
| ASV_263 | Proteobacteria; Alphaproteobacteria; Rhizobiales; Beijerinckiaceae; 1174-901-12 | 1.3091778 | 0.017303540 | HealyRT | enriched |
| ASV_285 | Acidobacteria; Acidobacteriia; Acidobacteriales; Acidobacteriaceae_(Subgroup_1); NA | 1.3017322 | 0.037551291 | HealyRT | enriched |
| ASV_294 | Proteobacteria; Alphaproteobacteria; Rhizobiales; Beijerinckiaceae; NA | 1.4061753 | 0.009596027 | HealyRT | enriched |
| ASV_443 | Proteobacteria; Alphaproteobacteria; Acetobacterales; Acetobacteraceae; NA | 1.1241770 | 0.047925505 | HealyRT | enriched |
| ASV_502 | Planctomycetes; Phycisphaerae; Tepidisphaerales; WD2101_soil_group; NA | 1.1595914 | 0.046870418 | HealyRT | enriched |
| ASV_1339 | Proteobacteria; Alphaproteobacteria; Acetobacterales; Acetobacteraceae; NA | 1.0947313 | 0.029295930 | HealyRT | enriched |
| ASV_2185 | Acidobacteria; Acidobacteriia; Acidobacteriales; Acidobacteriaceae_(Subgroup_1); Acidipila | 0.9197215 | 0.016223461 | HealyRT | enriched |
| ASV_77 | Actinobacteria; Actinobacteria; Frankiales; NA; NA | -1.0004512 | 0.012377682 | HealyRT | reduced |
| ASV_124 | Verrucomicrobia; Verrucomicrobiae; Chthoniobacterales; Chthoniobacteraceae; NA | -1.6683647 | 0.015719208 | HealyRT | reduced |
| ASV_135 | Proteobacteria; Alphaproteobacteria; Acetobacterales; Acetobacteraceae; NA | -1.0431617 | 0.015900713 | HealyRT | reduced |
| ASV_145 | Actinobacteria; Actinobacteria; Frankiales; Frankiaceae; Jatrophihabitans | -1.1406303 | 0.037176551 | HealyRT | reduced |
| ASV_160 | Planctomycetes; Phycisphaerae; Tepidisphaerales; WD2101_soil_group; NA | -1.1519487 | 0.015558312 | HealyRT | reduced |
| ASV_181 | Proteobacteria; Alphaproteobacteria; Acetobacterales; Acetobacteraceae; NA | -1.1178921 | 0.046192148 | HealyRT | reduced |
| ASV_329 | Actinobacteria; Acidimicrobiia; IMCC26256; NA; NA | -1.5619907 | 0.007276132 | HealyRT | reduced |
| ASV_345 | Proteobacteria; Alphaproteobacteria; Rhizobiales; Beijerinckiaceae; 1174-901-12 | -1.6905543 | 0.004453161 | HealyRT | reduced |
| ASV_424 | Proteobacteria; Alphaproteobacteria; Caulobacterales; Caulobacteraceae; PMMR1 | -1.3925957 | 0.032171082 | HealyRT | reduced |
| ASV_491 | Proteobacteria; Alphaproteobacteria; Acetobacterales; Acetobacteraceae; NA | -0.8992677 | 0.025750103 | HealyRT | reduced |
| ASV_1040 | Armatimonadetes; Armatimonadia; Armatimonadales; NA; NA | -0.8892736 | 0.043501351 | HealyRT | reduced |
| ASV_1119 | Actinobacteria; Actinobacteria; Frankiales; Acidothermaceae; Acidothermus | -0.9980928 | 0.014923945 | HealyRT | reduced |
